# Reversal of lactate and PD-1-mediated macrophage immunosuppression controls growth of PTEN/p53-deficient prostate cancer

**DOI:** 10.1101/2022.05.12.490223

**Authors:** Kiranj Chaudagar, Hanna M. Hieromnimon, Rimpi Khurana, Brian Labadie, Taghreed Hirz, Shenglin Mei, Raisa Hasan, Jordan Shafran, Anne Kelley, Eva Apostolov, Ghamdan Al-Eryani, Kate Harvey, Srikrishnan Rameshbabu, Mayme Loyd, Kaela Bynoe, Catherine Drovetsky, Ani Solanki, Erica Markiewicz, Marta Zamora, Xiaobing Fan, Stephan Schürer, Alex Swarbrick, David B. Sykes, Akash Patnaik

## Abstract

PTEN loss-of-function occurs in approximately 50% of mCRPC patients, and is associated with a poor prognosis, therapeutic outcomes and resistance to immune-checkpoint inhibitors. Recent clinical studies demonstrated that dual PI3K/AKT pathway inhibition and androgen axis blockade led to a modest improvement in progression-free survival of PTEN-deficient mCRPC patients, but the mechanistic basis for this limited efficacy is unknown. To elucidate potential resistance mechanism(s), we performed co-clinical trials in a prostate-specific PTEN/p53-deficient genetically-engineered mouse model, and discovered that the recruitment of PD-1-expressing tumor-associated macrophages (TAM) thwarts the phagocytosis-mediated anti-tumor efficacy of androgen deprivation therapy (ADT)/PI3K inhibitor (PI3Ki) combination. Strikingly, we observed a TAM-dependent ∼3-fold enhancement in the overall response rate with the addition of PD-1 antibody (aPD-1) to ADT/PI3Ki combination therapy. Mechanistically, decreased lactate production from PI3Ki-treated tumor cells suppressed histone lactylation (H3K18lac) within TAM, resulting in their phagocytic activation, which was augmented by concurrent ADT/aPD-1 treatment. Consistent with our murine observations, single cell RNA-sequencing analysis of human metastatic PC samples revealed a direct correlation between high glycolytic activity and phagocytosis suppression. Critically, feedback activation of Wnt/β-catenin signaling observed in non-responder mice following ADT/PI3Ki/aPD-1 combination treatment, restored lactate-mediated H3K18lac and suppressed phagocytosis within macrophages. Altogether, these data suggest that reversal of lactate and PD-1-mediated TAM immunosuppression by PI3Ki and aPD-1, respectively, controls tumor growth in combination with ADT, and warrants further clinical investigation in PTEN/p53-deficient mCRPC patients.

**One Sentence Summary:** Inhibition of tumor-cell intrinsic lactate production suppresses PTEN/p53-deficient prostate cancer growth via macrophage activation/phagocytosis

## INTRODUCTION

Prostate cancer (PC) is the second most commonly diagnosed cancer in men worldwide, with an estimated 375,304 deaths each year (*1*). While androgen deprivation therapy (ADT) remains the current standard of care for advanced PC, the majority of patients eventually progress and develop lethal metastatic castration-resistant PC (mCRPC) (*2*). Recent clinical trials have led to the FDA approval of androgen receptor signaling inhibitors (ARSI, abiraterone, enzalutamide, apalutamide) or docetaxel chemotherapy in combination with ADT for treatment of metastatic castration-sensitive PC (mCSPC) (*3–6*). Furthermore, recent studies have also shown a survival benefit of docetaxel chemotherapy in combination with AR antagonist darolutamide in mCSPC, relative to docetaxel alone (*7*). While the shift of ARSI and chemotherapy from the mCRPC to the mCSPC setting have resulted in an incremental improvement in overall survival, the majority of patients still progress to metastatic, castrate-resistant prostate cancer (mCRPC), which is associated with significant morbidity and mortality (*2, 4, 8, 9*). Furthermore, the use of chemohormonal therapies earlier in the disease continuum has led to the emergence of aggressive-variant prostate cancers (AVPC), that are associated with genomic loss-of-function (LOF) alterations in Phosphatase and tensin homolog (PTEN), TP53 and Retinoblastoma (Rb) genes and exhibit lineage plasticity and neuroendocrine/small cell histopathologic features (*10, 11*). Therefore, there is a critical unmet need to develop novel definitive therapies that eradicate AVPC.

Recently, there has been renewed interest in immunotherapy for the treatment of advanced PC, partly based on the anti-tumor immune activation that occurs with ADT, and partly based on the clinical responses to immune checkpoint inhibitors (ICI) targeting CTLA-4 and PD-1/PD-L1 in other cancers (*12–14*). However, only 10-25% of mCRPC patients respond to ICI, with a lack of durable benefit in the majority of patients (*15, 16*). Multiple mechanisms of immunotherapy resistance have been postulated in PC. These include a sparse immune infiltrate within the tumor microenvironment (TME), with a paucity of pre-existing T cells and a relative enrichment of innate immunosuppressive myeloid cells, such as tumor-associated macrophages (TAM) and myeloid derived suppressor cells (MDSC) (*17–21*). Furthermore, prior studies in PC have demonstrated that the release of immunosuppressive chemokines, such as TGF-β and IL-6, and the downregulation of MHC Class I expression correlates with reduced survival and increased distant metastasis, respectively (*22, 23*).

PTEN LOF alterations, which occur in approximately 50-75% of mCRPC patients, are associated with poor prognosis and development of metastases (*24–26*). In addition to enhanced tumor cell proliferation and survival via hyperactivation of the phosphatidylinsositol kinase-3 pathway (*27*), PTEN LOF alterations have been associated with an immunosuppressive tumor microenvironment, manifested by an increase in PD-L1 expression in some PTEN-deficient malignancies, increased expression of immunosuppressive chemokines such as CCL2, VEGF, IL1β, decreased infiltration and function of effector T cells, as well as the enhanced function of regulatory T cells (*18, 28, 29*). Co-occurrence of TP53-mutations is present in 56% of PTEN-deficient mCRPC patients, which also leads to an immunosuppressive TME by increasing TAM/MDSC infiltration (*26, 30*). Given the aggressive natural history and poor therapeutic outcomes of PTEN/p53 mutant advanced PC to standard-of-care hormonal therapies (*24*), chemotherapies (*31*) and ICI (*28, 32*), a deeper understanding of immune evasion mechanisms is critical for the discovery of new therapeutic strategies to effectively treat this molecular subset of AVPC.

One potential strategy to treat PTEN-deficient mCRPC is inhibition of the dysregulated activity of the PI3K signaling cascade, which drives increased aerobic glycolysis via the Warburg effect (*27*) and enhanced proliferation (*33*), survival (*34*), and metastasis (*25*). However, preclinical and clinical trial data has demonstrated that PI3K inhibitors are ineffective as single agents in the majority of solid tumor malignancies, independent of PTEN status (*35–38*). Furthermore, recent clinical studies have demonstrated that the combination of ADT, cytochrome (CYP) 17,20-lyase inhibitor abiraterone acetate and AKT inhibitor ipatasertib has shown modest improvement in progression-free survival (18.5 months in abiraterone/ipatasertib vs. 16.5 months in abiraterone/placebo) in PTEN-deficient mCRPC (*39*), but the mechanism of intrinsic and/or acquired resistance to this combinatorial approach remains unknown.

To elucidate the mechanistic basis for limited anti-cancer efficacy of dual PI3K/AKT axis and androgen blockade, we performed a pre-clinical trial of degarelix (ADT), with and without copanlisib (pan-PI3K inhibitor) in the castrate-resistant prostate-specific PTEN/p53-deficient genetically engineered mouse model (GEMM) (*40*), which recapitulated the modest anti-tumor responses observed in mCRPC patients (*39*). Importantly, we discovered that the recruitment of PD-1 expressing tumor-associated macrophages (TAM) limits the phagocytosis mediated-anti-tumor efficacy elicited by ADT/copanlisib combination treatment. We therefore tested the hypothesis that the addition of a PD-1 blocking antibody (aPD-1) to the ADT/copanlisib combination could enhance the anti-tumor efficacy in GEMMs, and observed an ∼3-fold enhancement of overall response rate (ORR), which was abrogated *in vivo* by TAM depletion. Mechanistically, decreased lactate production from copanlisib treated tumor cells resulted in suppression of histone lactylation (H3K18lac) within TAM, resulting in their phagocytic activation which was augmented by concurrent ADT/aPD-1. Critically, feedback activation of Wnt/β-catenin signaling observed in non-responder mice following ADT/PI3Ki/aPD-1 combination treatment, restored lactate-mediated H3K18lac and suppressed phagocytosis within macrophages. Taken together, our findings demonstrate that reversal of lactate and PD-1-mediated TAM immunosuppression by PI3Ki and aPD-1, respectively, in combination with ADT controls tumor growth and warrants further clinical investigation in PTEN/p53-deficient mCRPC.

## RESULTS

### ADT/PI3Ki combination inhibits tumor growth in a PTEN/p53-deficient murine PC model in a tumor cell-extrinsic manner

Prior clinical studies have demonstrated limited ADT responsiveness in patients with PTEN-deficient AVPC (*24*). As a first step towards elucidating the combinatorial impact of ADT with PI3Ki in PTEN/p53-deficient murine PC, 16-20 week old Pb-Cre; PTEN^fl/fl^ Trp53^fl/fl^ mice bearing established 100-150 mm^3^ solid tumors were treated with degarelix (Luteinizing hormone-releasing hormone (LHRH)-antagonist, chemical castration), singly and in combination with copanlisib (pan-PI3K inhibitor) for 4 weeks. Tumor growth was monitored using non-invasive magnetic resonance imaging (MRI). While ADT significantly decreased serum testosterone levels in all mice to castrate levels (*41*), the ORR was 16.7%, demonstrating that the majority of mice were *de novo* resistant to ADT **(Fig. 1A and fig. S1A-C)**. Copanlisib, either as a single-agent or in combination with ADT, completely inhibited the PI3K pathway and demonstrated an increased ORR of 37.5% and 25%, respectively, relative to untreated control **(Fig. 1A and fig. S2A-C)**, thus corroborating published clinical trial data showing modest improvement in PFS with dual targeting of PI3K/AKT-pathway axis and androgen blockade, relative to androgen blockade alone (*42*). In particular, a partial response (PR)/tumor shrinkage was observed with degarelix/copanlisib combination treatment (12.5%) relative to single agents (0%), suggesting a synergistic interaction. Interestingly, copanlisib treatment with or without androgen depleted media (AD, which mimics ADT *in vivo*) had no effect *in vitro* on cell proliferation and survival in PTEN/p53-deficient murine prostate GEMM tumor derived cell lines from 4-month (adenocarcinoma, AC1, (*43*)) and 7-month old (sarcomatoid, SC1, (*43*)) mice, respectively **(fig. S3A-C)**. In contrast, tumor extracts harvested after 7 days of ADT/copanlisib combination treatment revealed a 2.3-fold decrease in Ki67+ tumor cells (a marker of proliferation measured by flow cytometry) in the TME relative to single agents and untreated controls **(Fig. 1B)**. These findings suggest that ADT/PI3Ki combination exerts its anti-cancer activity via a tumor cell-extrinsic mechanism in PTEN/p53-deficient prostate GEMM.

**Fig. 1.**
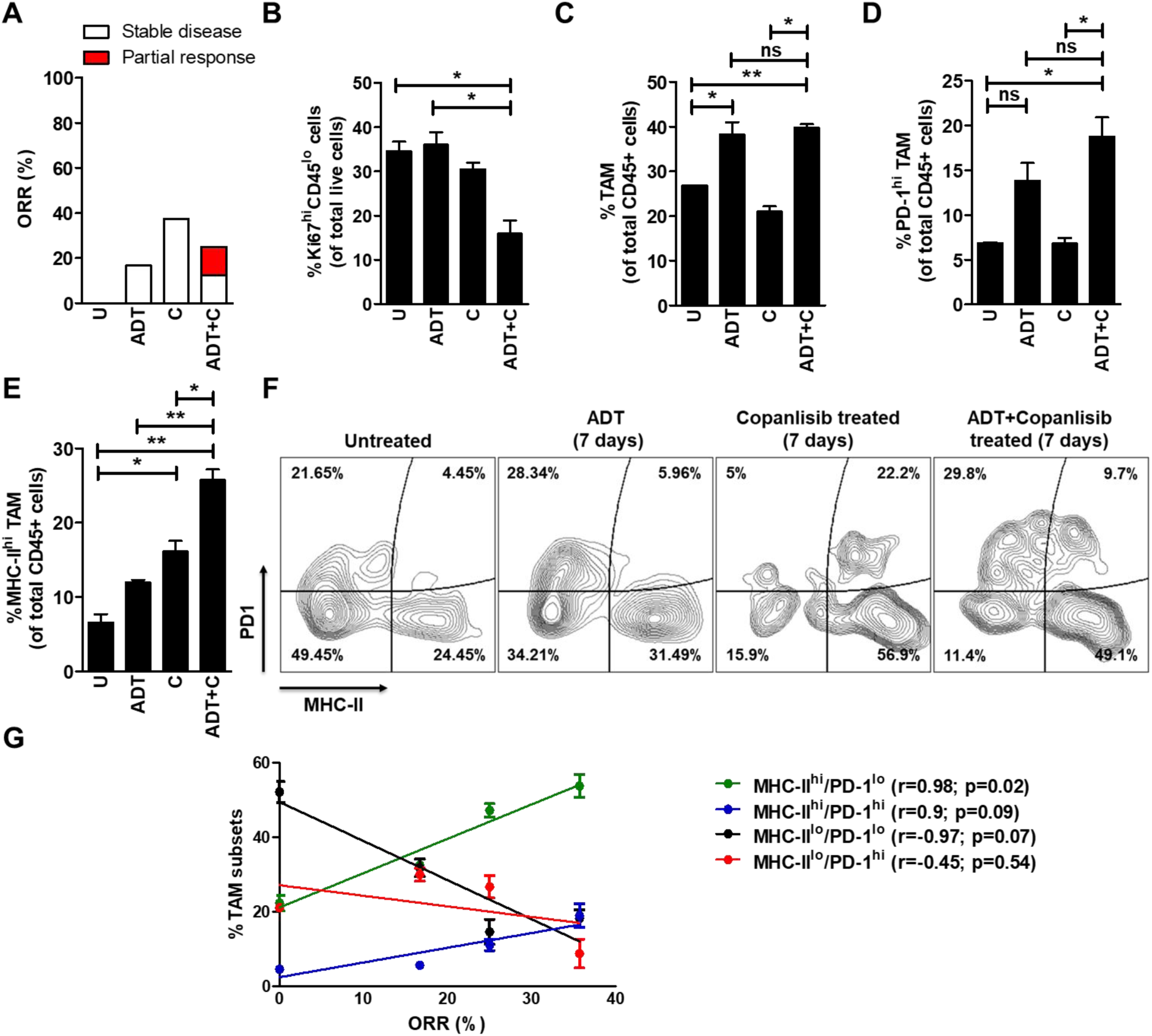
The combination of ADT and PI3Ki demonstrates partial anti-tumor response in PTEN/p53-deficient murine PC via increased infiltration/activation of tumor-associated macrophages (TAM) within the TME. **(A)** Pb-Cre; PTEN^fl/fl^ Trp53^fl/fl^ mice were followed by serial ultrasound until they developed solid prostate tumors (100-150 mm^3^) by 16-20 weeks of age. Mice were randomized to untreated, degarelix (0.625 mg, single dose, ADT) treated, copanlisib (14 mg/kg, *iv*, every alternate day, PI3Ki, C) treated, and their combination treated groups. Tumor growth was monitored using MRI, and ORR (partial response + stable disease) were calculated, as described in Methods. (**B-F**) Following acute treatment for 7 days, prostate tumors were harvested and analyzed by flow cytometry for the following cell populations: proliferating tumor cells (Ki67^+^CD45^-^, (**B**)), total TAM (CD45^+^CD11b^+^F4/80^+^ cells, (**C**)), PD-1 (**D**) and MHC-II (**E**) expressing TAM. (**F**) Representative flow plots demonstrate % frequency of MHC-II^hi/lo^/PD-1^hi/lo^ expressing TAM (CD45^+^CD11b^+^F4/80^+^ cells). (**G**) Correlation plot highlights association of MHC-II^hi^/PD-1^lo^, MHC-II^hi^/PD-1^hi^, MHC-II^lo^/PD-1^lo^ and MHC-II^lo^/PD-1^hi^ TAM subsets with % ORR observed following 28-days treatment of single agents or their combination and in untreated group (U). n=2-8 mice per group. Significances/p-values were calculated by Chi-square test (panel A), one-way ANOVA (panel B-E), Pearson test (panel G) and indicates as follows, *p<0.05 and **p<0.01; ns = not statistically significant.

### ADT/PI3Ki combination increases frequency of activated macrophages in PTEN/p53- deficient murine PC

To elucidate the anti-cancer mechanism for ADT/PI3Ki combination within the TME, we performed unbiased immune profiling of prostate tumors derived from Pb-Cre; PTEN^fl/fl^ Trp53^fl/fl^ mice following 7 days of treatment with degarelix, singly and in combination with copanlisib. The TME of untreated PTEN/p53-deficient prostate tumors were enriched with tumor associated-macrophages (TAM, 25% of total CD45^+^ cells) and granulocytic-myeloid derived suppressive cells (Gr-MDSC, 38% of total CD45^+^ cells), with a relative paucity of T cells (CD4-expressing, 3% and CD8-expressing, 1% of total CD45^+^ cells) **(Fig. 1C and fig. S4A-C**). We observed a 1.4- fold increase in the frequency of total TAM in PTEN/p53-deficient tumor bearing mice treated with degarelix, relative to untreated control, which was not enhanced with concurrent copanlisib **(Fig. 1C)**. Furthermore, degarelix/copanlisib treatment led to a significant increase of PD-1 expressing TAM, relative to untreated controls and copanlisib alone **(Fig. 1D)**. Importantly, copanlisib treatment led to a 2.3-fold increase in frequency of activated (MHC-II^hi^) TAM, which was significantly enhanced to 3.6-fold in combination with ADT, relative to untreated control **(Fig. 1E)**. In contrast, either ADT or copanlisib, singly and in combination showed a 1.6-fold decrease in the frequency of Gr-MDSCs and a 2.8/5-fold increase in CD4^+^/CD8^+^T cells within the TME, which was not enhanced with the combination **(fig. S4A-C**). Taken together, these data demonstrate that the combination of ADT/PI3Ki significantly enhances TAM activation within TME, relative to corresponding single-agents.

As ADT/PI3Ki combination increased frequency of both MHC-II and PD-1 expressing TAM relative to untreated control we focused on the role of MHC-II^hi/lo^ and PD-1^hi/lo^ TAM subsets in tumor growth control and found that activated TAM (MHC-II^hi^/PD-1^lo^ and MHC-II^hi^/PD-1^hi^) were positively correlated with ORR (r=0.98 and 0.9, respectively), whereas a negative association was observed between immunosuppressive MHC-II^lo^/PD-1^lo^ and MHC-II^lo^/PD-1^hi^ TAM and ORR (r=-0.97 and -0.45, respectively, **Fig. 1F-G**). Furthermore, the positive correlation was significant with MHC-II^hi^/PD-1^lo^TAM (p=0.02), but not with MHC-II^hi^/PD-1^hi^TAM (p=0.09, **Fig. 1G**). Collectively, these data demonstrate that TAM activation maximally correlates with an anti-tumor response, which is attenuated by PD-1 expression within TAM.

### AD/PI3Ki combination enhances phagocytosis of PTEN/p53-deficient murine prostate tumor cells via activated TAM

As a first step towards elucidating a mechanism by which activated (MHC-II^hi^/PD-1^lo^ and MHC-II^hi^/PD-1^hi^) TAM control tumor growth in Pb-Cre; PTEN^fl/fl^ Trp53^fl/fl^ mice following degarelix/copanlisib treatment, we examined the effects of copanlisib in combination with androgen depletion (AD, mimicking *in vivo* degarelix treatment) on TAM-mediated phagocytosis of PTEN/p53-deficient GEMM-tumor derived PC cell lines, AC1 and SC1. We first performed co-culture experiments to determine the relative phagocytic activity of sorted TAM subsets (MHC-II^hi/lo^ and PD-1^hi/lo^ TAM) co-cultured with AC1/SC1 cells at baseline **(Fig. 2A)**. Consistent with the TAM subset/ORR correlation data **(Fig. 1G)**, the activated MHC-II^hi^/PD-1^lo^ TAM exhibited the highest phagocytic activity, with a 14.5-fold increase in uptake of AC1/SC1 cells relative to the MHC-II^lo^/PD-1^hi^ TAM. In contrast, the MHC-II^hi^/PD-1^hi^ and MHC-II^lo^/PD-1^lo^ TAM populations demonstrated an 8.5-fold and 2.5-fold increase in phagocytosis of AC1/SC1 cells, relative to the MHC-II^lo^/PD-1^hi^ TAM (**Fig. 2B)**. Importantly, activated (MHC-II^hi^) TAM had significantly lower levels of PD-L1 and CD206 expression relative to the inactivated (MHC-II^lo^) TAM population, independent of PD-1 status. Furthermore, CD206 expression was higher in inactivated MHC-II^lo^/PD-1^hi^ TAM, relative to their inactivated MHC-II^lo^/PD-1^lo^ counterparts, when co-cultured with either AC1 or SC1 cells **(Fig. 2C and D)**. Collectively, these data demonstrate that PD-1 expression within TAM enhances their immunosuppressive phenotype and suppresses their phagocytic activity, independent of their activation status.

**Fig. 2.**
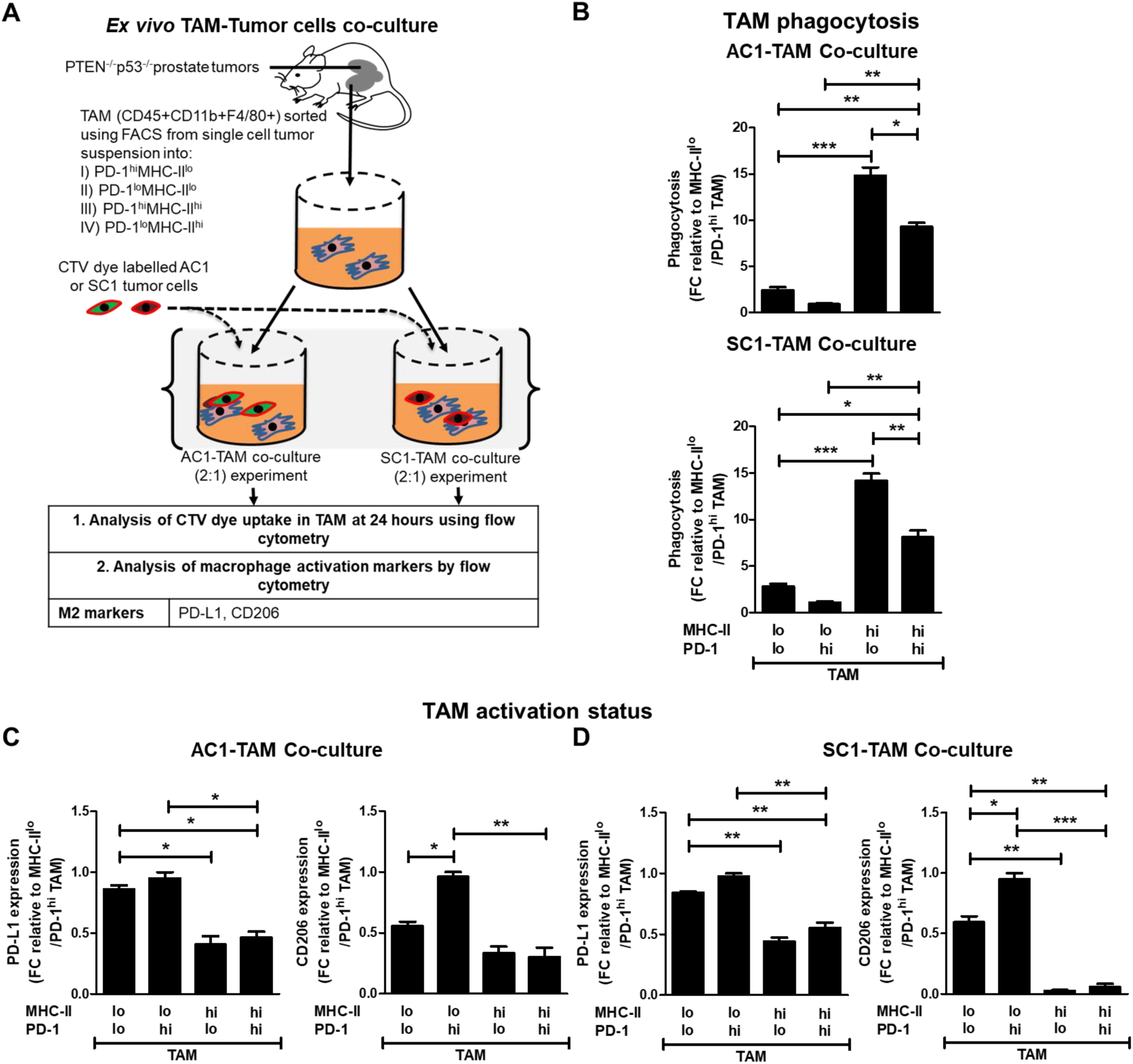
PD-1 upregulation suppresses phagocytic capacity of activated TAM. **(A)** Experimental schema for phagocytosis experiment, in which sorted TAM from PTEN/p53-deficient prostate GEMM tumors are co-cultured with CTV dye stained-AC1/SC1 cells for 24 hours. **(B)** Bar graphs demonstrate fold change (FC) in phagocytosis of AC1 (upper panel) and SC1 (lower panel) cells by MHC-II^hi/lo^/ PD-1^hi/lo^ expressing TAM, relative to untreated group. PD-L1 and CD206 expression were analyzed in the TAM subsets following co-culture with AC1 **(C)** and SC1 **(D)** cells. n=2 independent experiments. Significances/p-values were calculated by one-way ANOVA and indicates as follows, *p<0.05, **p<0.01 and ***p<0.001.

Next, we examined the impact of AD, singly and in combination with copanlisib on phagocytic uptake of AC1 and SC1 cells *ex vivo*, across the 4 TAM subsets described above. Whereas AD resulted in a striking increase in phagocytosis of AC1 and SC1 cells by MHC-II^hi^/PD-1^lo^ TAM (19.6-fold and 3.8-fold, respectively), relative to untreated controls, it did not alter the phagocytic activity of MHC-II^hi^/PD-1^hi^ TAM relative to untreated group (**Fig. 3B and C),** suggesting a suppressive effect of PD-1 expression on ADT-induced phagocytosis. In contrast, single-agent copanlisib treatment showed an increase in phagocytosis of AC1 and SC1 cells by both MHC-II^hi^/PD-1^lo^ TAM (11-fold and 3-fold, respectively) and MHC-II^hi^/PD-1^hi^ TAM (12-fold and 7-fold, respectively), relative to untreated controls. Importantly, AD/copanlisib combination resulted in a 32-fold and 9-fold increase in phagocytosis of AC1 and SC1 cells, respectively, by MHC-II^hi^/PD-1^lo^ TAM, relative to untreated controls. The combination also resulted in a 20.4-fold and 11.8-fold increase in phagocytosis of AC1 and SC1 cells, respectively, by MHC-II^hi^/PD-1^hi^ TAM, relative to untreated controls **(Fig. 3B and C)**. Neither AD/copanlisib combination or corresponding single agent treatments altered phagocytic activity of MHC-II^lo^/PD-1^lo^ or MHC-II^lo^/PD-1^hi^ TAM, relative to untreated controls **(Fig. 3B and C)**. Collectively, these data demonstrate that AD and PI3Ki have differential effects on activated TAM subsets and their phagocytosis of PTEN/p53-deficient prostate tumor cells, resulting in enhanced effects with the combination relative to single-agents. Furthermore, AD/PI3Ki mediated phagocytosis is significantly suppressed in inactivated TAM subsets.

**Fig. 3.**
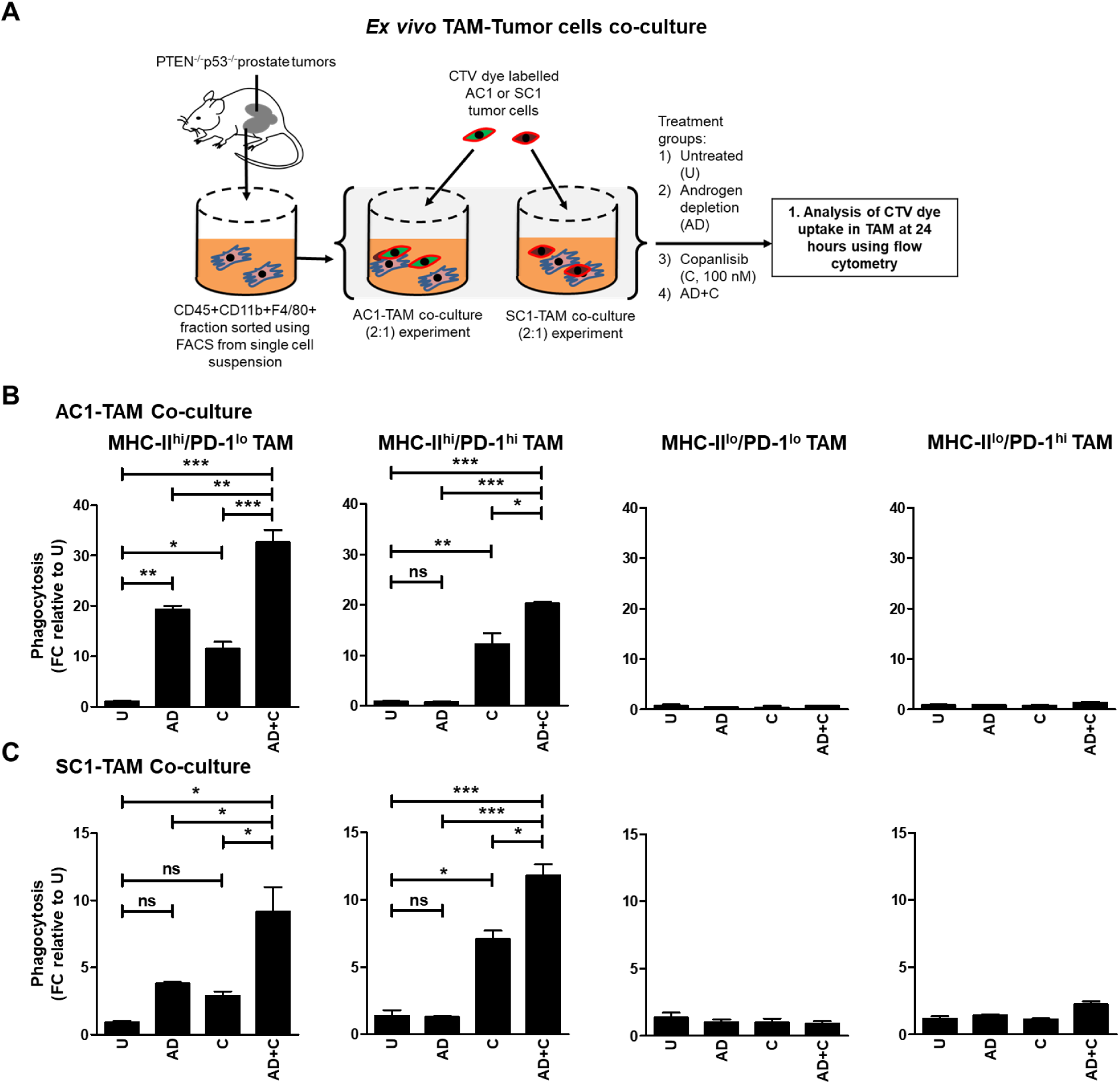
ADT/PI3Ki combination therapy significantly enhances phagocytosis of PTEN/p53- deficient murine cancer cells by activated TAM. **(A)** Experimental schema for phagocytosis experiment, in which sorted TAM from PTEN/p53-deficient prostate GEMM tumors were co-cultured with CTV dye stained-AC1/SC1 cells under normal and AD conditions, and treated with either DMSO (negative control) or copanlisib (C, 100 nM) for 24 hours. Bar graphs demonstrate fold change (FC) in phagocytosis of AC1 **(B)** and SC1 **(C)** cells by MHC-II^hi^/PD-1^lo^, MHC-II^hi^/PD-1^hi^, MHC-II^lo^/PD-1^lo^ and MHC-II^lo^/PD-1^hi^ TAM subsets, relative to untreated group (U). n=2 independent experiments. Significances/p-values were calculated by one-way ANOVA and indicates as follows, *p<0.05, **p<0.01 and ***p<0.001; ns = not statistically significant.

### The addition of PD-1 blockade to AD/PI3Ki enhances phagocytic capacity of suppressive PD-1^hi^ macrophages

Given our findings that baseline and AD/PI3Ki-induced phagocytosis of PTEN/p53- deficient GEMM tumor-derived cells is attenuated by high PD-1 expression in the activated TAM subset **(Fig. 2B & 3B)**, we next evaluated the impact of aPD-1 on TAM-mediated phagocytosis of AC1/SC1 cells either alone or in combination with copanlisib, with or without AD **(Fig. 4A)**. We observed a significant increase in the phagocytic activity of fluorescence-activated cell sorting (FACS)-isolated suppressive MHC-II^lo^/PD-1^hi^ TAM towards AC1/SC1 cells following treatment with AD/copanlisib/aPD-1 combination, relative to AD/copanlisib combination (2.7- and 2.1-fold, respectively), aPD-1 monotherapy (9.3- and 3.9-fold, respectively) and untreated controls (28- and 11.7-fold, respectively, **Fig. 4B and C)**. This was accompanied by an increase in MHC-II expression on suppressive MHC-II^lo^/PD-1^hi^ TAM following AD/copanlisib/aPD-1 combination treatment, relative to untreated control, indicating TAM reprogramming by AD/copanlisib/aPD-1 combination **(fig. S5A and B)**. Given the higher baseline phagocytic activity of the MHC-II^hi^/PD- 1^hi^ TAM subpopulation, the relative increase with aPD-1 addition is more modest relative to the MHC-II^lo^/PD-1^hi^ subpopulation, as demonstrated by the AD/copanlisib/aPD-1 combination showing a partial increase in phagocytosis of AC1/SC1 cells, relative to AD/copanlisib combination (1.4 and 1.1-fold, respectively), aPD-1 monotherapy (12.8- and 4.8-fold, respectively) and untreated controls (32- and 12-fold, respectively, **Fig. 4B and C**). As anticipated, the addition of aPD-1 did not alter the phagocytic capacity of MHC-II^hi^/PD-1^lo^ and MHC-II^lo^/PD- 1^lo^ TAM subpopulations, relative to their corresponding controls without aPD-1 **(fig. S6A and B)**. Consistently, AD/copanlisib combination increased MHC-II expression in the MHC-II^lo^/PD-1^lo^ subset relative to the untreated, but this was not accentuated by addition of aPD-1 **(fig. S5A and B)**. These data suggest that blockade of PD-1 in the suppressive PD-1^hi^ TAM overcomes the immunosuppressive effects of PD-1 on phagocytosis of tumor cells within the PTEN/p53-deficient TME.

**Fig. 4.**
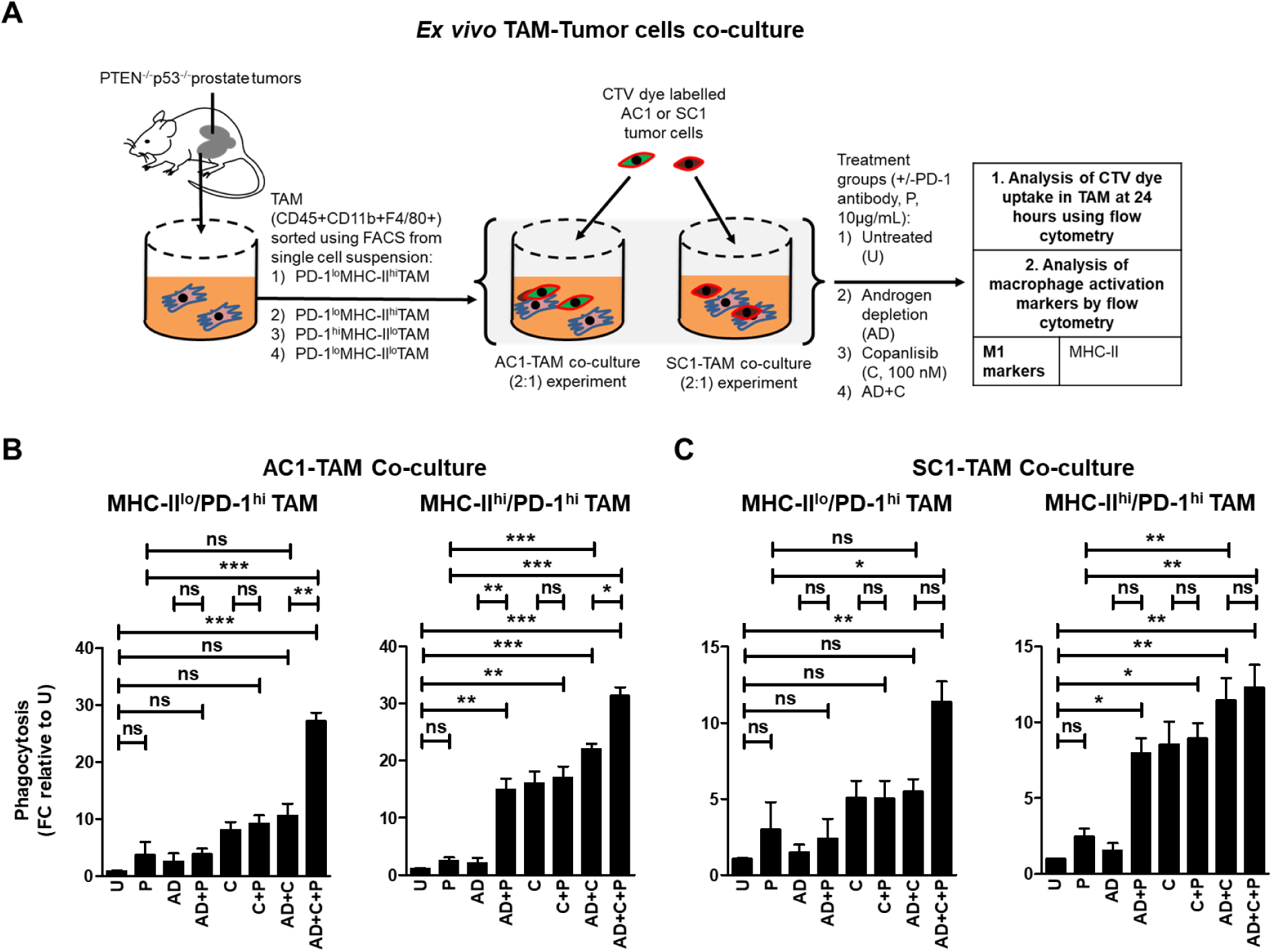
The addition of PD-1 blockade to androgen depletion/PI3Ki enhances phagocytic capacity of suppressive PD-1^hi^ macrophages. **(A)** Schema illustrating co-culture of FACS-sorted TAM subsets (MHC-II^hi/lo^/ PD-1^hi^) from PTEN/p53-deficient prostate tumors with CTV dye stained-AC1/SC1 cells under normal and AD conditions. Co-cultures were treated with copanlisib (C, 100 nM), PD-1 antibody (P, 10 μg/mL) or their combination for 24 hours, followed by phagocytosis assay measuring CTV dye uptake within TAM. Bar graphs demonstrate fold change (FC) in phagocytic activity of MHC-II^hi/lo^/PD-1^hi^ expressing TAM, relative to untreated group (U) in AC1 **(B)** and SC1 **(C)** cells. n=2 independent experiments. Significances/p-values were calculated by one-way ANOVA and indicates as follows, *p<0.05, **p<0.01 and ***p<0.001; ns = not statistically significant.

The addition of aPD-1 in combination with AD resulted in a 7.4- and 5.7-fold increase in the phagocytosis of AC1 and SC1 cells, respectively, by MHC-II^hi^/PD-1^hi^ TAM subset, relative to AD controls. Interestingly, AD/aPD-1 combination was insufficient to enhance phagocytosis in the MHC-II^lo^/PD-1^hi^ TAM, relative to AD alone, likely related to the higher threshold of activation in this immunosuppressive TAM subset **(Fig. 4B and C)**. As expected, the AD/aPD-1 did not enhance phagocytosis of the MHC-II^hi^/PD-1^lo^ and MHC-II^lo^/PD-1^lo^ TAM subset, relative to AD alone group **(fig. S6A and B)**. Copanlisib/aPD-1 did not increase phagocytic activity of MHC-II^hi^/PD-1^hi^ and MHC-II^lo^/PD-1^hi^ TAM towards AC1 or SC1 cells, relative to copanlisib monotherapy **(Fig. 4B and C)**. Taken together, these data demonstrate that the addition of aPD-1 to AD or AD/copanlisib combination (not copanlisib alone) enhances phagocytic activity of PD-1^hi^ activated TAM.

We also analyzed CD47 and PD-1 expression by flow cytometry to explore the impact of AD/copanlisib/aPD-1 combination treatment on the expression of phagocytosis inhibitory checkpoints on tumor cells (*44*). None of the treatment groups had an impact on expression of these checkpoints on AC1/SC1 cells **(fig. S7A and B)**. Collectively, these data demonstrate that concurrent AD/copanlisib/aPD-1 blockade significantly enhances both activation and phagocytosis of MHC-II^hi^ TAM when co-cultured with PTEN/p53-deficient tumor cells, relative to corresponding singlet and doublet treatments.

### Direct treatment of activated TAM with AD, not PI3Ki, enhances phagocytic capacity

To understand how AD impacts the functionality of TAM, we pre-treated FACS-sorted TAM with AD, PI3Ki or their combination and then performed phagocytosis experiments using PTEN/p53-deficient GEMM tumor-derived AC1 and SC1 cells as target populations **(Fig. 5A)**. Interestingly, AD demonstrated a 6.5-fold increase in the phagocytic capacity of MHC-II^hi^/PD-1^lo^ TAM, relative to untreated control, which was not enhanced by concurrent copanlisib and/or aPD-1 blockade. Critically, MHC-II^hi^/PD-1^hi^ TAM required the addition of aPD-1 to achieve a similar induction of phagocytosis **(Fig. 5B and C)**. AD/aPD-1 combination did not alter phagocytic activity of MHC-II^lo^/PD-1^lo^ TAM and MHC-II^lo^/PD-1^hi^ TAM **(fig. S8A and B)**. Consistent with the observed phagocytic data **(Fig. 5B and Fig. 5C)**, AD activated TAM directly, which was not accentuated with concurrent copanlisib or aPD-1 **(fig. S9A and B)**. Taken together, these data demonstrate that AD facilitates an antitumor immune response in PTEN/p53-deficient PC by increasing total TAM infiltration and MHC-II^hi^/PD-1^lo^ TAM subset activation/phagocytic activity, with enhanced phagocytosis of MHC-II^hi^/PD-1^hi^ activated TAM subset requiring concurrent PD-1 blockade.

**Fig. 5.**
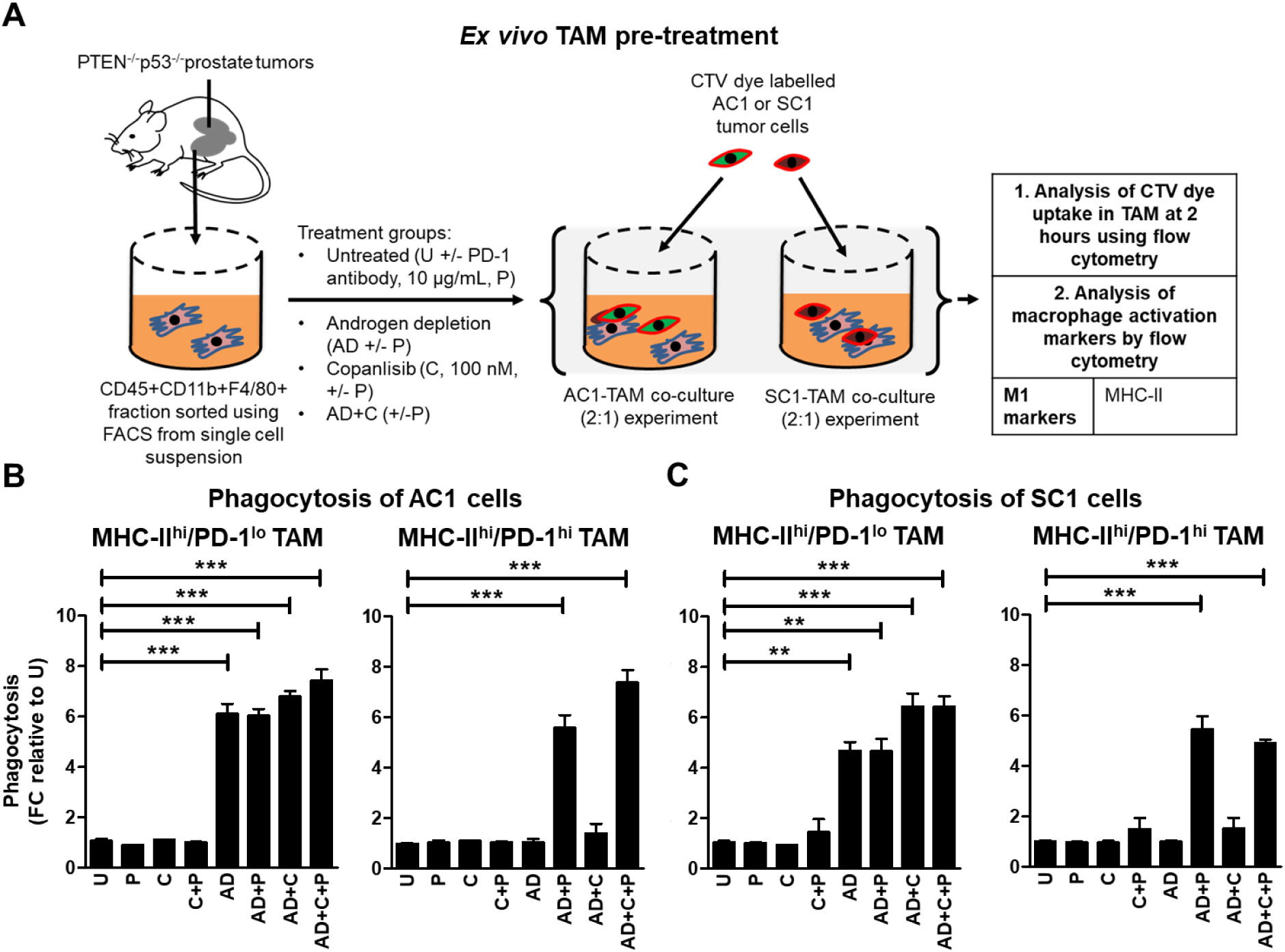
Pre-treatment of TAM *ex vivo* with androgen depletion alone and in combination with aPD-1, increases phagocytosis of PTEN/p53-deficient GEMM tumor cells by MHC-II^hi^/PD-1^lo^ and MHC-II^hi^/PD-1^hi^ TAM subsets, respectively. **(A)** Schema illustrating pre-treatment of FACS-sorted TAM subsets from PTEN/p53-deficient prostate tumors with copanlisib (C, 100 nM), PD-1 antibody (P, 10 μg/mL) or their combination under normal and AD condition for 24 hours. After PBS wash, treated TAM were co-cultured with CTV dye stained-AC1/SC1 cells for 2 hours. Bar graphs demonstrate fold change (FC) in phagocytosis of AC1 **(B)** and SC1 **(C)** cells by MHC-II^hi^/PD-1^hi/lo^ expressing TAM, relative to untreated group (U). n=2 independent experiments. Significances/p-values were calculated by one-way ANOVA and indicates as follows, **p<0.01 and ***p<0.001; ns = not statistically significant.

### PI3Ki enhances phagocytic capacity of activated TAM by inhibiting histone lactylation

As direct treatment with copanlisib did not *directly* alter the activation status and phagocytic capacity of activated TAM **(Fig. 5B-C and fig. S9A-B)**, we hypothesized that copanlisib contributes to anti-cancer immunity by overcoming the immunosuppressive secretome of PTEN/p53-deficient PC cells, and *indirectly* enhancing TAM activation/phagocytosis. To test this hypothesis, we performed *ex vivo* conditioned media (CM) experiments **(Fig. 6A)**, and observed a 4.5-fold increase in the phagocytic capacity of activated MHC-II^hi^/PD-1^lo^ and MHC-II^hi^/PD-1^hi^ TAM **(Fig. 6B)**, but not inactivated MHC-II^lo^/PD-1^lo^ and MHC-II^lo^/PD-1^hi^ TAM, in response to CM collected from single cell suspensions of tumors following treatment with copanlisib **(fig. S10A and B)**. Since the PI3K pathway is a pivotal driver of glucose metabolism and lactate production by cancer cells (*45*), we next assessed the impact of copanlisib on lactate release in the ex vivo CM. We observed a 18.1% decrease in lactate levels in response to copanlisib, relative to untreated controls **(fig. S11)**. Corroborating this finding, we observed a similar decrease in lactate production from AC1/SC1 cells in response to copanlisib **(fig. S11)**. Given recent findings that histone lactylation (lactylation of lysine 18 on histone-3, H3K18lac) can drive immunosuppression within TAM (*46*), we tested the hypothesis that a decrease in lactate content in the *ex vivo* CM was accompanied by reduced histone lactylation within activated TAM. Importantly, we observed a significant decrease in H3K18lac specifically with MHC-II^hi^/PD-1^lo^ and MHC-II^hi^/PD-1^hi^ TAM subsets **(fig. 6C)**, suggesting that PI3Ki-induced lactate suppression within tumor cells and reduced histone lactylation within activated TAM enhances phagocytosis of tumor cells. To specifically address this hypothesis, we tested the impact of exogenous lactate addition to the *ex vivo* CM on phagocytic capacity of activated TAM **(Fig. 6A)**. Interestingly, we observed an abrogation of copanlisib-induced enhanced phagocytosis of AC1/SC1 cells by activated MHC-II^hi^/PD-1^lo^ and MHC-II^hi^/PD-1^hi^ TAM **(Fig. 6B)**, which was accompanied by restoration of histone lactylation within these TAM subsets **(Fig. 6C)**. Importantly, direct treatment with copanlisib did not alter the histone lactylation status of TAM **(fig. S12)**. Collectively, these data demonstrate that copanlisib treatment decreases lactate production from treated PTEN/p53- deficient PC cells and secondary histone lactylation within MHC-II^+^ TAM subsets, resulting in enhanced TAM activation/phagocytosis within the TME.

**Fig. 6.**
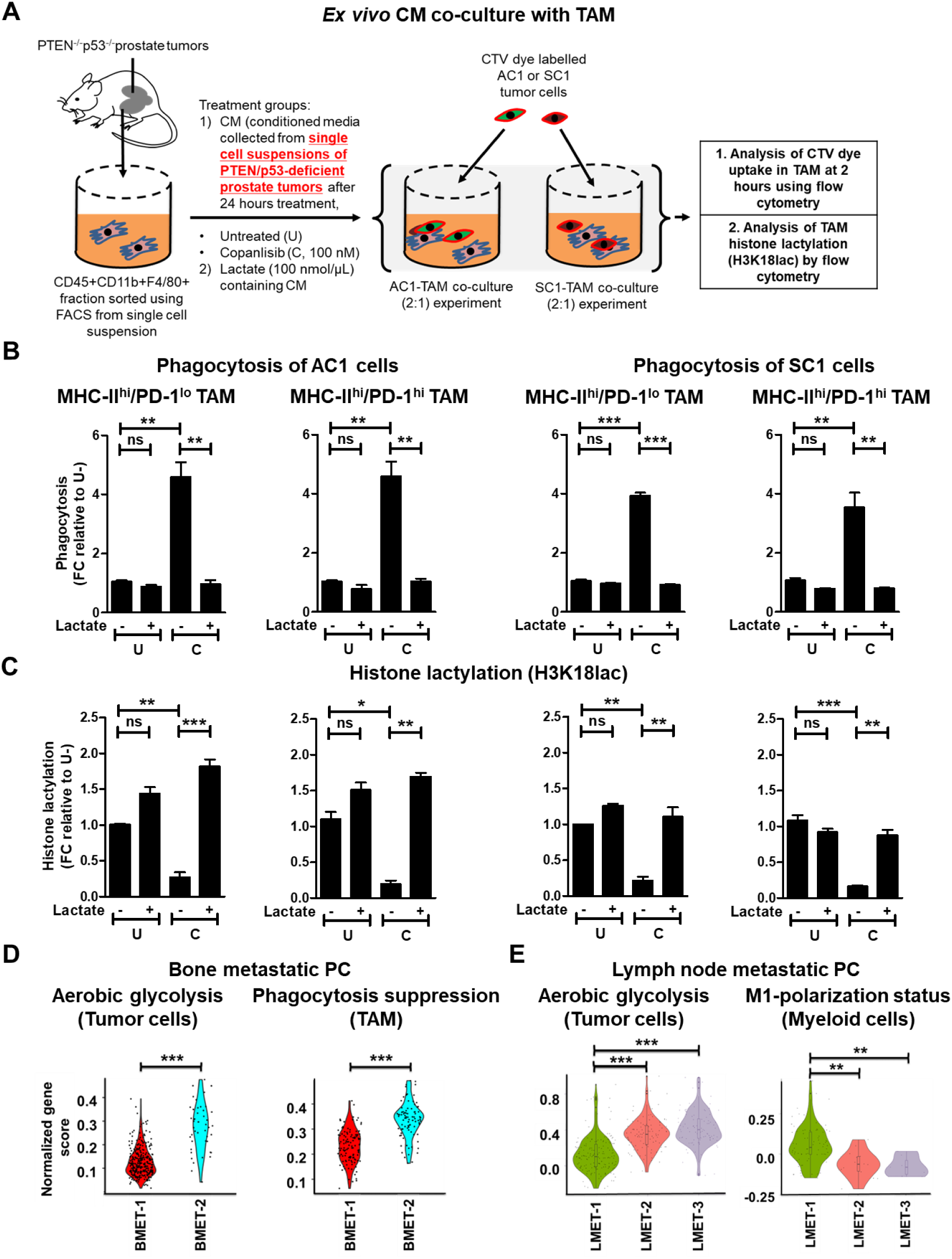
PI3Ki suppresses lactate production from treated PTEN/p53-deficient PC cells and secondary histone lactylation within activated TAM, resulting in enhanced TAM phagocytosis. **(A)** Schema illustrating lactate add-back phagocytosis experiment on *ex vivo* CM. Single cell suspensions of PTEN/p53-deficient prostate GEMM tumors were treated with copanlisib (C, 100 nM) and CM was collected 24 hours following treatment. FACS-sorted TAM from PTEN/p53-deficient prostate tumors were cultured *ex vivo* with CM for 24 hours in presence or absence of lactate (100 nmol/μL). After PBS wash, TAM were co-cultured with CTV dye stained-AC1/SC1 cells for 2 hours. Bar graphs demonstrate fold change (FC) in phagocytic activity **(B)** and histone lactylation status **(C)** of activated MHC-II^hi^/PD-1^lo^ and MHC-II^hi^/PD-1^hi^ TAM, relative to untreated group (U). **(D-E)** Single cell suspensions were prepared from human bone/lymph node metastatic PC patient samples (BMET-1/-2 and LMET-1/-2/-3) and underwent single-cell RNA using Chromium controller 10x Genomics platform. Annotated tumor cells, TAM and myeloid cells were characterized for aerobic glycolysis **(D, E)**, phagocytosis suppression **(D)** and M1-polarization status **(E)**, respectively, using a normalized gene score, as described in the methods section. For *ex vivo* studies, n=2 independent experiments and for bioinformatics analysis, n = 355 tumor cells and 389 TAM in bone metastatic PC patients; n = 367 tumor cells and 101 myeloid cells in lymph node metastatic PC patients. Significances/p-values were calculated by one-way ANOVA (panel B-C), Wilcoxon rank-sum text (panel D-E) and indicates as follows, *p<0.05, **p<0.01 and ***p<0.001; ns = not statistically significant.

To validate our preclinical findings demonstrating a mechanistic link between aerobic glycolytic activity i.e. lactate production from tumor cells, and macrophage phagocytosis/activation in advanced human PC, we performed single cell RNA sequencing (scRNAseq) of bone metastatic PC patient samples, BMET-1 and BMET-2. We observed significantly higher aerobic glycolytic activity in cancer cells and a correspondingly higher suppression of TAM phagocytosis in BMET-2 vs BMET-1 **(Fig. 6D)**. In parallel, we also conducted scRNA-seq in mCRPC patients metastatic lymph node samples (LMET-1, -2 and -3). While the number of TAM were insufficient to perform a phagocytosis gene signature analysis, we observed an inverse correlation between aerobic glycolytic activity within tumor cells and M1-polarization status within myeloid cells **(Fig. 6E)**, thus validating a role for tumor cell intrinsic glycolysis in evading TAM-mediated innate immune phagocytic response within the metastatic PC microenvironment.

### ADT/PI3Ki/aPD-1-induced macrophage activation *in vivo* controls tumor growth in 60% of Pb-Cre; PTEN^fl/fl^ Trp53^fl/fl^ mice

Based on our findings that ADT, aPD-1 and PI3Ki treatment can overcome TAM immunosuppression via distinct mechanisms across the 4 TAM subsets, we determined whether ADT/PI3Ki/aPD-1 combination enhances anti-cancer responses in Pb-Cre; PTEN^fl/fl^ Trp53^fl/fl^ mice, relative to singlet and doublet controls. We treated established prostate tumor-bearing mice with aPD-1, singly or in combination with copanlisib and/or ADT, and measured the tumor growth kinetics using MRI. The addition of aPD-1 to ADT increased the ORR to 33.3% (compared to 16.7% with ADT alone, **fig. S1B**) at 28 days **(Fig. 7A and fig. S13A-B).** No increase in ORR was observed with the addition of aPD-1 to copanlisib (37.5%), relative to copanlisib alone (37.5%) **(Fig. 7A and fig. S13C),** which mirrors our co-culture phagocytosis data **(Fig. 4B and C)**. Importantly, mice treated with ADT/copanlisib/aPD-1 combination demonstrated a significantly increased ORR of 60% by 28 days, relative to singlet and doublet controls **(Fig. 7A, 1A and fig. S13A-D)**, and correlated with our *ex vivo* phagocytosis data **(Fig. 4B and C)**. This ORR was accompanied by an increase in frequency of activated TAM in the TME **(Fig. 7B)**. Critically, clodronate co-treatment, which depletes activated (MHC-II^hi^/PD-1^lo^ and MHC-II^hi^/PD-1^hi^) TAM, completely abolished tumor control induced by ADT/copanlisib/aPD-1 combination **(Fig. 7A-B and fig. S14A-C)**. Additionally, cytokine array analysis demonstrated an increase in pro-inflammatory mediators (IL-1α, TNFα, CCL-22, CCL-5, IL-6 and CXCL-16) and a decrease in anti-inflammatory cytokines (M-CSF and CCL-6) from single cell suspensions of tumors following treatment with ADT + copanlisib + aPD-1 **(table S1)**, which supports our primary hypothesis that activated/repolarized macrophages are largely responsible for the observed anti-tumor response elicited by the triple combination. Collectively, these data demonstrate that the triple therapeutic combination (ADT + PI3Ki + aPD-1) drives TAM activation and substantially increased tumor control *in vivo* in PTEN/p53-deficient GEMM mice, relative to the ADT/PI3Ki combination.

**Fig. 7.**
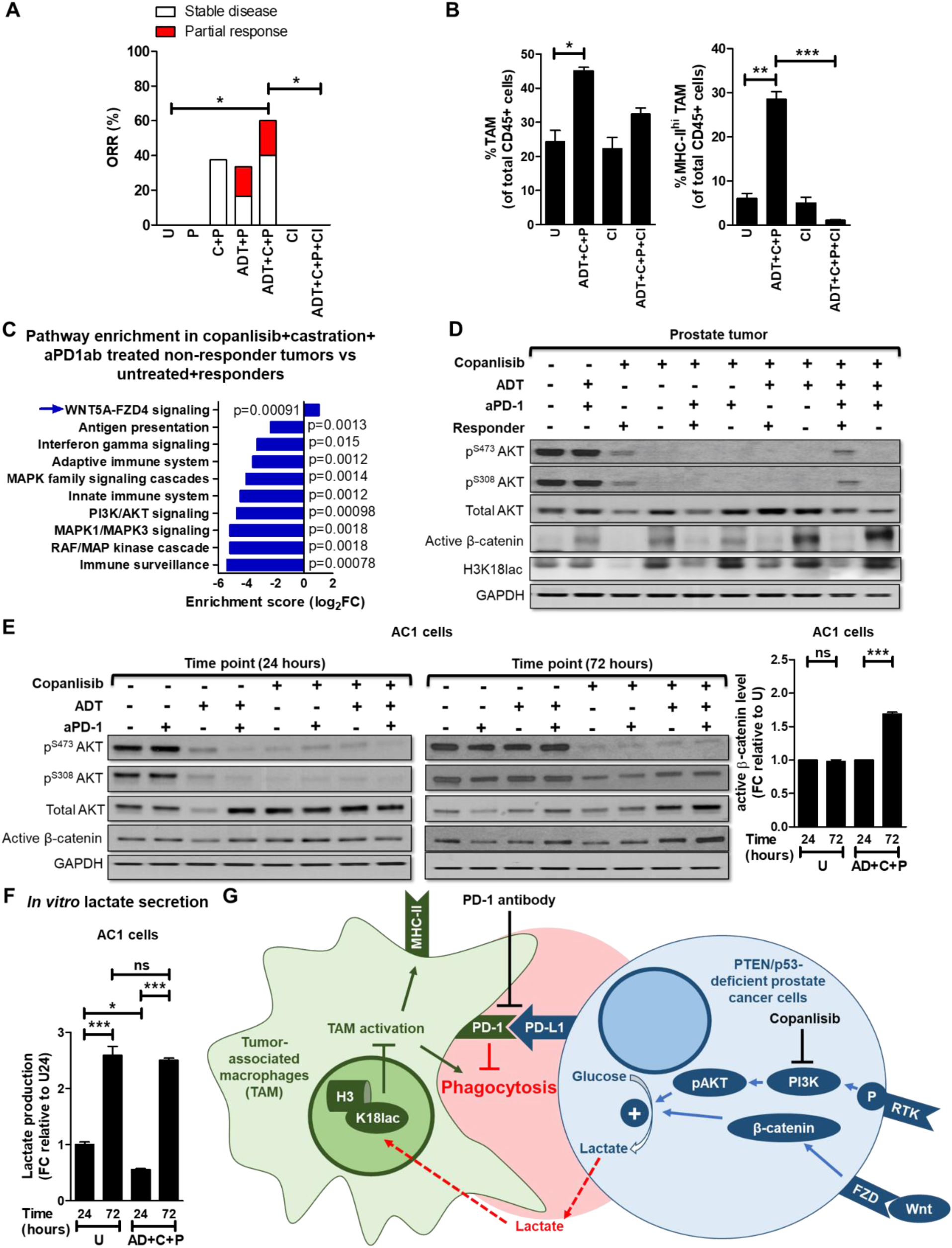
ADT + PI3Ki + aPD-1 antibody leads to tumor control in PTEN/p53-deficient GEMM via activated TAM, with feedback Wnt/β-catenin activation mediated restoration of lactate and histone lactylation observed in non-responders. **(A)** Pb-Cre; PTEN^fl/fl^ Trp53^fl/fl^ mice with established prostate tumors were treated with PD-1 antibody (aPD-1, 100 μg/mouse, *ip*, every alternate day, P) alone or in combination with ADT (degarelix, 0.625mg, single dose, castration) or copanlisib, (14 mg/kg, *iv*, every alternate day, C) or ADT + copanlisib. For *in vivo* macrophage depletion, clodronate (200 μg/mouse, *ip*, every week, Cl) was administered concurrently with ADT + copanlisib + PD-1 antibody. Tumor volume was monitored non-invasively using MRI and ORR (partial response + stable disease) was calculated, as described in Methods. **(B)** Pb-Cre; PTEN^fl/fl^ Trp53^fl/fl^ mice were treated with the indicated drug(s) for 7 days, and analyzed for % frequency of total and MHC-II expressing TAM (CD45^+^CD11b^+^F4/80^+^ cells). **(C)** Tumor RNA was isolated following 28 days treatment with indicated agents and sequenced to perform pathway enrichment analysis. Reactome plot shows pathway analysis for differentially expressed genes in non-responder tumors, relative to responder and untreated. **(D)** Western blot analyses were performed for indicated protein markers on PTEN/p53-deficient prostate GEMM tumor extracts following treatment with ADT, copanlisib, PD-1 antibody or their combinations for 28 days. **(E-F)** AC1 cells were treated with copanlisib (C, 100 nM), PD-1 antibody (P, 10 μg/mL) or their combinations under normal or AD conditions for 24 and 72 hours. Western blot analyses **(E)** were performed for indicated protein markers on AC1 cell lysates, and active β-catenin level was quantified by Image J software, and lactate levels in the supernatant were analyzed by fluorimetry. **(F)**. **(G)** Model illustrating the decrease in lactate production from PTEN/p53-deficient PC cells in response to PI3Ki inhibitor treatment overcomes histone lactylation mediated TAM suppression and enhances phagocytosis of PC cells within TAM. Concurrent ADT and aPD-1 treatment *directly* activates TAM within TME, thereby accentuating the effects of single-agent PI3Ki on TAM functionality/polarization. Wnt/β-catenin activation drives resistance to ADT/PI3Ki/aPD-1 combination therapy. For *in vivo* studies, n=6-10 mice per group (panel A), n=2 mice per group (panel B), and for *in vitro*, n=3 independent experiments. Significances/p-values were calculated by Chi-square test (panel A), one-way ANOVA (panel B and F), Reactome analysis (panel C) and indicated as follows, *p<0.05, **p<0.01 and ***p<0.001; ns = not statistically significant.

### Feedback activation of Wnt/β-catenin signaling observed in non-responder Pb-Cre; PTEN^fl/fl^ Trp53^fl/fl^ mice following ADT/PI3Ki/aPD-1 combination treatment, restores lactate-mediated histone lactylation (H3K18lac) and suppresses macrophage phagocytosis

To elucidate the mechanism of resistance to ADT/PI3Ki/aPD-1 combination therapy in the non-responder mice, RNAseq analysis was performed on tumors from non-responder mice and compared to responder and untreated mice. Pathway enrichment analysis revealed upregulation of the Wnt pathway and down-regulation of immune responses in non-responder tumors, relative to untreated and responder tumors **(Fig. 7C)**. Consistent with this data, Western blot analysis on tumor extracts demonstrated an increase in active β-catenin expression and restoration of H3K18lac in non-responder tumors relative to responder mice across all treatment groups **(Fig. 7D)**. *In vitro* western blot analysis of PTEN/p53-deficient tumor-derived AC1/SC1 cells demonstrated a concomitant increase in active β-catenin expression in AC1/SC1 cells following copanlisib + aPD-1 treatment under AD conditions for 72 hours, relative to 24-hour treatment. This feedback activation of Wnt/β-catenin signaling was accompanied by an increase in lactate production at 72 hours, following an initial decrease with acute copanlisb/AD/aPD-1 combination treatment *in vitro* **(Fig. 7E-F and fig. S15A-C)**. Critically, treatment of BMDM with *in vitro* CM collected at 72 hours following AD/copanlisib/aPD-1 combination treatment demonstrated an abrogation of phagocytic activity and an increase in H3K18lac, relative to 24 hours CM treatment. **(fig. S16A-B)**. Taken together, these data suggest that Wnt-β-catenin pathway activation and restoration of tumor cell/TAM cross-talk via lactate secretion and histone lactylation drive resistance to ADT/PI3Ki/aPD-1 combination therapy **(Fig. 7G)**.

## DISCUSSION

Immune-based therapies have failed to provide a meaningful benefit in the majority of mCRPC patients (*15*), underscoring the critical need to develop rational IO combination strategy to effectively treat and eradicate mCRPC. PTEN LOF occurs in approximately 50% of mCRPC patients, and is associated with a poor prognosis, immunosuppressive TME and resistance to ICI (*26, 30, 47*). Hyperactivated PI3K signaling in the setting of PTEN LOF enhances glucose consumption within tumor cells by increasing GLUT-1/4 on the plasma membrane and activating hexokinase 2, which results in a metabolic switch towards aerobic glycolysis and increased lactate secretion (*27*). Lactate can promote M2/MHC-II^lo^-TAM polarization by inhibiting vacuolar ATPase mediated HIF2α stabilization and promoting epigenetic reprograming via histone lactylation (*46, 48, 49*), and resultant immune evasion (*50*). Furthermore, lactate has been shown to have differential metabolic roles within the TAM subpopulations in the TME. For example, lactate diminishes glycolytic activity in M1/MHC-II^hi^ TAMs, but upregulates the TCA cycle, enhances Arg1, NOS2 expressions in MHC-II^lo^ TAMs, resulting in suppression of T cell proliferation (*51*). We observed TAM phagocytosis suppression and decreased myeloid M1 signatures with corresponding high glycolytic activity in bone and lymph node metastatic PC patients, respectively, validating our mechanistic findings in advanced PC patients. Critically, we discovered that PI3Ki combination therapy induces MHC-II^hi^ TAM activation/phagocytosis by selectively decreasing lactate production from PTEN/p53-deficient PC cells and resultant histone lactylation, particularly on H3 at lysine 18 within MHC-II^hi^/PD-1^lo^ and MHC-II^hi^/PD-1^hi^ TAM, but not in MHC-II^lo^/PD-1^lo^ TAM and MHC-II^lo^/PD-1^hi^ TAM. Concurrent ADT and aPD-1 treatment overcomes immunosuppression *directly* within MHC-II^hi^/PD-1^lo^ TAM and MHC-II^lo^/PD-1^hi^ TAM, respectively, resulting in the reinvigoration of anti-tumor macrophage-mediated phagocytic response and sensitization of PTEN/p53 murine PC to ICI. Collectively, these findings demonstrate that therapeutic suppression of the Warburg effect within tumor cells, when combined with *direct* macrophage activation strategies, can significantly overcome TAM-mediated immunosuppression within the TME of PTEN/p53-deficient AVPC.

As noted above, the generation of TAM-driven anti-cancer innate immune phagocytic responses to ADT/PI3Ki/aPD-1 combination treatment were largely driven by the MHC-II^hi^/PD-1^hi/lo^ TAM and MHC-II^lo^/PD-1^hi^ subpopulations. In contrast, the MHC-II^lo^/PD-1^lo^ TAM were not reprogrammed by ADT, aPD-1 blockade or PI3Ki-mediated lactate suppression and were unable to phagocytose tumor cells, thereby contributing to therapy resistance. To overcome this limitation, the blockade of additional phagocytic checkpoints could be explored. For example, phagocytosis is negatively regulated by the SIRPα/CD47 (*52*), MHC-I/LILRB1 (*53*) and CD24/Siglec-10 axes (*54*). We found CD47 expression on PTEN/p53-deficient PC cells, which has been previously shown to interact with SIRPα of MHC-II^lo^ TAM as well as MHC-II^hi^ TAM (*55*), thereby activating SHP-1/-2 and inhibiting phagocytosis (*44, 52*). In addition, we observed Wnt/β-catenin pathway activation as a potential resistance mechanism to ADT/PI3Ki/aPD-1 combination therapy, which has been previously shown to upregulate CD47 expression on cancer cells by increasing cMyc-transcriptional activity (*56*). Our findings provide a mechanistic rationale for exploring anti-CD47 antibody in combination with ADT/PI3Ki/aPD-1 to further enhance macrophage-driven anti-cancer responses in aggressive-variant PTEN/p53-deficient PC. Furthermore, our co-culture phagocytosis assay showed a higher phagocytosis of PTEN/p53-deficient GEMM tumor-derived AC1 (adenocarcinoma) cells, relative to SC1 (sarcomatoid) cells in MHC-II^hi^/PD-1^lo^ TAM in response to AD/PI3Ki+/-aPD-1 combination treatment, underscoring the need to perform high-content CRISPR screening on AC1, SC1 tumor cells or MHC-II^lo^/PD-1^lo^ TAM (*57*), which could shed light on additional unidentified phagocytosis checkpoints and their contribution to treatment resistance.

Given prior studies demonstrating a reciprocal feedback relationship between AR and PI3K signaling in PTEN-deficient PC, current clinical trials are co-targeting the PI3K/AKT-pathway and AR-blockade in PTEN-deficient mCRPC patients (*37, 38, 42, 58, 59*). Ipatasertib (a pan-AKTi), in combination with abiraterone (IPATential-150 trial) has demonstrated a modest improvement in progression-free survival (rPFS) of PTEN-loss mCRPC patients (*39*). Furthermore, capivasertib (a pan-AKT inhibitor)/abiraterone combination is under clinical investigation for PTEN-deficient metastatic hormone sensitive PC (NCT04493853, CAPItello-281 trial). Our preclinical mechanistic studies demonstrate that ADT plus PI3Ki drives tumor cell extrinsic TAM-mediated tumor growth control in 25% of PTEN/p53-deficient GEMM mice through MHC-II^hi^ TAM activation and phagocytosis of cancer cells. Furthermore, we discovered that responses were thwarted by PD-1 expressing TAM population which was partly overcome by the concurrent addition of aPD-1, resulting in a 60% ORR. Based on our pre-clinical findings, we predict that infiltration of PD-1 expressing TAMs following dual AR and PI3K/AKT axis blockade will be a key acquired resistance mechanism to the combinatorial approach in IPATential-150 (*39*) and CAPItello-281 trials (NCT04493853), thus warranting further clinical/translational investigation of concurrent PI3K/AKT axis and PD-1 blockade with ADT in PTEN-deficient hormone-sensitive and mCRPC setting.

Our findings suggest that Wnt/β-catenin pathway activation drives resistance to PI3Ki- based therapy in PTEN/p53-deficient murine PC via restoration of lactate production from tumor cells and resultant TAM suppression. Prior studies have demonstrated that APC-mutant colon cancer generates increased lactate production from tumor cells via upregulation of Wnt pathway (*60*). Furthermore, PI3Ki treatment has been shown to increase transcription of Wnt-ligands and phosphorylation of the co-receptor LRP5/6, which dismantles the β-catenin degradation complex and results in Wnt/β-catenin pathway activation mediated resistance in colorectal cancer (*61*). These findings highlight the cross-talk between oncogenic signaling pathways to preserve the Warburg effect, which contributes to the development of immunometabolic resistance when the PI3K pathway is targeted with precision medicine therapies. Taken together, our results suggest that addition of Wnt/β-catenin pathway inhibitor, such as a Porcupine inhibitor, in combination with ADT/PI3Ki/PD-1 blockade therapy can potentially overcome resistance, and warrants further investigation in combination with ADT/PI3Ki/aPD-1 triplet therapy to treat PTEN/p53-deficient AVPC.

## MATERIALS AND METHODS (table S2 contains detail on chemical and reagents)

### *In vivo* murine treatment and prostate tumor growth kinetic studies

Experiments were performed in accordance with NIH guidelines and protocol approved by the Institutional Animal Care and Use Committee (IACUC) at University of Chicago. Prostate-specific PTEN/p53- deficient (Pb-Cre; PTEN^fl/fl^ Trp53^fl/fl^) mice were screened for autochthonous prostate tumor development at 16 weeks of age by ultrasound. Following the development of solid tumors (when the solid tumors reached a long-axis diameter of 5 mm under ultrasound imaging), the mice were treated with either degarelix (0.625 mg/mouse, *sc*, every 28-days, ADT group), copanlisib (14mg/kg, *iv*, every alternate day), PD-1 antibody (aPD-1, 200 μg/mouse, *ip*, every alternate day) single agents or their combinations (as indicated in Figure legends). Tumors were monitored at baseline and every 14-days following treatment using 9.4 Tesla small animal MRI scanner (Bruker, Germany) with 11.6 cm inner diameter and actively shielded gradient coils (maximum constant gradient strength = 230mT/m for all axes). Amira software was used to outline region of interest and delineate cystic and solid disease components on MRI. A Matlab script summed over the pixels (1 pixel = 5μm^3^) in these outlined regions was utilized to compute tumor volume. Solid prostate tumor volume was derived by subtracting cystic volume from total prostate tumor volume. Mice were considered treatment responsive if solid tumor volume did not grow >20% (stable disease, SD) at indicated time point following treatment, relative to baseline. Furthermore, % Partial response (PR) was calculated based on a proportion of total mice that exhibited >30% decrease in solid tumor volume following treatment, relative to baseline. % Overall Response Rate (ORR) was calculated based on number of responder mice (SD + PR), relative to total mice enrolled in a specific treatment.

### Serum testosterone determination

To demonstrate that chemical castration with degarelix generates castrate levels of testosterone, Pb-Cre; PTEN^fl/fl^ Trp53^fl/fl^ mice were treated with degarelix (0.625 mg, *sc*, single dose, 28 days, ADT group) and blood was drawn from tail vein at baseline, 3 and 28-days following treatment. Serum was isolated and analyzed for testosterone level as per manufacturer ELISA protocol.

### Cell lines and culture conditions

For *in vitro* experiments on tumor cells, PTEN/p53-deficient prostate GEMM tumor derived cancer cell lines, AC1 (adenocarcinoma type) and SC1 (sarcomatoid type) were cultured in prostate epithelial growth media (PrEGM) in the absence (AC1 medium) and presence (SC1 medium) of 10% fetal bovine serum (FBS), respectively (*62*). PrEGM was generated by adding necessary supplements to Prostate Epithelial Basal Media (PrEBM), as per manufacturer’s protocol. To mimic *in vivo* ADT condition, AD-PrEGM was generated from PrEBM by adding all supplements, except hydrocortisone, a known androgen receptor agonist (*63*). This was labeled as AD-AC1 medium, whereas AD-SC1 medium had 10% charcoal stripped FBS (CSS) in AD-PrEGM.

### Western blot analysis

For *in vivo* studies, established prostate tumors from Pb-Cre; PTEN^fl/fl^ Trp53^fl/fl^ mice were harvested following treatment, lysed in T-PER buffer and 10 μg of total protein extracts for each sample underwent sodium dodecyl sulfate-polyacrylamide gel electrophoresis (SDS-PAGE), followed by Western Blot analysis. For *in vitro* studies, AC1 or SC1 cells were treated with indicated drug(s) under normal AC1/SC1 or AD-AC1/SC1 media conditions, respectively. Cells were lysed in RIPA buffer and 10 μg of total protein extracts underwent SDS-PAGE followed by Western blotting, and probed for pAKT-S473, pAKT-T308, total AKT, active-β-catenin, H3K18lac, pLRP6-S1490, LRP6 or GAPDH as indicated in Figure legends. Bands were quantified using Image J software.

### Flow cytometric analysis

Pb-Cre; PTEN^fl/fl^ Trp53^fl/fl^ mice were treated with the indicated drug(s), and harvested tumors were digested using liberase, and filtered through a 70 μm cell strainer. The resulting cell suspension was centrifuged and incubated for 2 minutes in 1 mL of Ammonium-Chloride-Potassium (ACK) solution to lyse red blood cells. ACK was neutralized with 10 mL of PBS and the cell suspension was centrifuged at 1800 rpm for 5 minutes. The cell pellet was resuspended at density of 1x10^6^ cells/mL in PBS containing either myeloid or lymphoid antibody cocktails. Myeloid antibody cocktails were prepared by resuspending 10 μL of anti-CD45, CD11b, CD11c, Ly6c, Ly6g, MHC-II, F4/80, PD-1, PD-L1 antibodies and 1 μL of Ghost-viability dye in 1 mL PBS. Lymphoid antibody cocktails were prepared by resuspending 10 μL of anti-CD45, CD3, CD4, CD8, CD19, FoxP3, Ki67, PD-1, PD-L1 antibodies and 1 μL of Ghost-viability dye in 1 mL PBS. After 30 minutes of incubation, stained single cell suspensions were washed with PBS and incubated with fixation buffer for 15 minutes. Following fixation, single cell suspensions of samples incubated with myeloid antibody cocktail were washed and resuspended in PBS for flow cytometry run whereas lymphoid antibody cocktail-stained cells were incubated overnight with 1 mL fix/perm buffer containing 10 μL of anti-FoxP3 and Ki67 antibodies for intracellular stains. Following completion of staining, cells were washed, resuspended in PBS and flow cytometry was performed for lymphoid markers. Flow cytometry data were gated for myeloid and lymphoid subsets and their activation status using FlowJo v 10.7 software.

### Proliferation and apoptosis assays

AC1/SC1 cells were treated with copanlisib (100 nM), aPD-1 (10 μg/mL) and their combinations in presence of AC1/SC1 or AD-AC1/SC1 media for indicated time points. Adherent cells were digested with trypsin and total number of cells per well counted using hemocytometer. Proliferation rate was calculated by dividing the number of cells at a given time point with the seeding density. For apoptosis assay, tumor cell suspensions were further stained with anti-Annexin V antibodies and propidium iodide (PI, DNA marker dye) as per protocol supplied with kit. Flow cytometry analysis was performed for annexin V+/PI- (apoptotic) and annexin V+/PI+ (necrotic) cells. % cell death was defined as the sum total of both apoptotic and necrotic cells frequencies.

### TAM isolation

To elucidate functional relevance of TAM and their activation states within the TME following treatment, established prostate tumors of GEMM mice were harvested, and single cell suspensions were prepared using liberase digestion method. Following ACK treatment, single cell suspensions were stained with anti-CD45, CD11b, F4/80 antibodies (1:100 dilution for each) and Ghost-viability dye (1:1000 dilution) in 1 mL SC1 media. TAM (CD45^+^CD11b^+^F4/80^+^ cells) were sorted using FACS. Furthermore, MHC-II and PD-1 antibodies (1:100 dilution for each) were added in staining cocktails to isolate the following TAM subsets using FACS: MHC-II^hi^/PD-1^lo^, MHC-II^hi^/PD-1^hi^, MHC-II^lo^/PD-1^lo^, and MHC-II^lo^/PD-1^hi^.

### CM collection

Established prostate tumors from untreated Pb-Cre; PTEN^fl/fl^ Trp53^fl/fl^ mice were harvested, and single cell suspensions were prepared using liberase digestion method. Cells were plated at density of 3x10^6^ cells per P100-culture dish. These suspensions were treated *ex vivo* with copanlisib (100 nM), aPD-1 (10 μg/mL) and their combinations under AC1 or AD-AC1 media conditions as indicated in Figure legends. Supernatants were collected following treatment, named as an *ex vivo* CM and utilized for proteomic, metabolic and TAM functionality experiments. For *in vitro* CM, supernatants were collected following treatment of AC1/SC1 cells with copanlisib (100 nM), aPD-1 (10 μg/mL) and their combinations in presence of AC1/SC1 or AD-AC1/SC1 media, as described in Figure legends.

### BMDM generation

To perform *in vitro* macrophage functionality assay, bone marrow progenitors were extracted from femur of male Cre^-/-^ GEMM mice by flushing twice with cold PBS buffer. Progenitors were cultured for 7 days in M-CSF (30 ng/mL) containing DMEM media supplemented with 10% FBS, 1% non-essential amino acid and 1% penicillin/streptomycin, to enable differentiation into bone marrow derived macrophages (BMDM, (*51*)), which were utilized for downstream functional studies.

### *Ex vivo* phagocytosis assay

TAM or their subsets were sorted from untreated PTEN/p53- deficient established prostate tumors using FACS and utilized for three distinct phagocytosis assay approaches. (i) AC1 and SC1 cells were stained with CTV dye (1:2000 dilution in PBS) and <u>co-cultured</u> with TAM subsets at a 2:1 ratio of tumor cells:TAM. These co-cultures were treated *ex vivo* with copanlisib (100 nM), aPD-1 (10 μg/mL) or their combinations for 24 hours in AC1/SC1 and AD-AC1/AD-SC1 media, to mimic conditions with and without androgen, respectively. (ii) TAM were either <u>pre-treated *ex vivo* directly</u> with copanlisib (100 nM), aPD-1 (10 μg/mL), or their combinations in presence of normal AC1/SC1 and AD-AC1/SC1 media or (iii) <u>indirectly with CM</u> (as described in above subsection) for 24 hours to dissect mechanism of TAM activation/phagocytosis. To investigate role of lactate on macrophage suppression, FACS-sorted TAM or BMDM were pre-treated for 24 hours with *ex vivo* or *in vitro* CM, in the presence or absence of lactate, which was added at a final concentration of either 50 or 100 nmol/μL. After these pre-treatments, TAM were co-cultured with CTV dye stained AC1/SC1 cells at Tumor cells:TAM ratio of 2:1 for 2 hours. At the end of phagocytosis, cells were then washed twice with PBS and stained with anti-CD45, MHC-II (M1-anti-tumor/activated macrophage marker), PD-1, (1:100 dilution for each), H3K18lac antibodies (1:100 dilution for each) and Ghost-viability dye (1:1000 dilution) in 1 mL PBS. Phagocytic activity and histone lactylation status of each TAM subset (MHC-II^hi^/PD-1^lo^, MHC-II^hi^/PD-1^hi^, MHC-II^lo^/PD-1^lo^, and MHC-II^lo^/PD-1^hi^ TAM) was calculated by normalizing MFI of CTV dye and H3K18lac staining, relative to untreated groups, respectively. Baseline phagocytic activity of TAM subsets was normalized relative to untreated MHC-II^lo^PD-1^hi^ TAM.

### *In vitro* phagocytosis checkpoint analysis

AC1/SC1 cells were treated with copanlisib (100 nM), aPD-1 (10 μg/mL) and their combinations in normal AC1/SC1 and AD-AC1/SC1 media conditions for 24 hours, to mimic conditions with and without androgen, respectively. Following trypsinization, cells were stained with anti-CD47 and PD-1 antibodies, and analyzed by flow cytometry.

### Lactate determination

To dissect mechanism of anti-tumor response with indicated drug combinations, quantification of lactate was performed using colorimetry kit on both *ex vivo* and *in vitro* CM, as described above.

### Single cell RNA sequencing and bioinformatic analysis

Bone metastatic PC samples were collected and handled in accordance to the protocol approved by the Institutional Review Board (Dana Farber/Harvard Cancer Center protocol 13-416 and Partners protocol 2017P000635/PHS). Following consent, spine metastatic PC patients underwent tumor specimen extraction procedure as described previously (*64*). Tumor specimens were enzymatically dissociated to single cells, treated with ACK lysis buffer to remove erythrocytes and suspended in Media 199 supplemented with 2% FBS. Single cells suspensions were stained with CD235 antibody and DAPI-dye. Cells that appeared negative for both stains were sorted using BD FACS Aria III instrument equipped with a 100um nozzle (BD Biosciences, San Jose, CA) and further sequenced for RNA by Chromium Controller as per manufacturer protocol (10x Genomics). FASTQ files were processed using CellRanger and human genome-hg19 was considered as the reference genome to generate the matrix files containing cell barcodes and transcript counts. Cells with total UMI exceeding 600 were further included in the downstream analysis. Quality control and data exploration were done by Pagoda2. Cells were annotated using Conos tutorial. A gene set signature score was used to measure cell states in different cell subsets and conditions. Signature score were calculated as average expression values of genes in a given set. Specifically, normalized gene signature score of each cell were calculated as an average normalized (for cell size) gene expression magnitude for; 1) aerobic glycolysis activity (ALDOA, BSG, ENO1, ENO2, ENO3, GPI, HK1, LDHA, LDHB, PFKL, PFKM, PFKP, PGAM1, PGAM2, PGK1, PKLR, PKM, SLC16A1, SLC16A3, SLC2A1, TPI1; (*65*)) in tumor cells and 2) non-phagocytic suppressive phenotypes (ATAD1, B2M, FCGR1, IGHM, IL4, MFF, STAP1, SYNE1; (*66*)) of TAM.

For metastatic lymph nodes of PC patients, baseline biopsies were collected and processed as mentioned in the investigator-initiated, IRB-approved clinical trial (University of Chicago, NCT03572478) of rucaparib in combination with nivolumab, co-sponsored by Clovis Oncology and Bristol Myers Squibb. Briefly, single cell suspensions were prepared in RPMI-1640 supplemented with 10% FBS, followed by enzymatic digestion and incubation in ACK lysis buffer. RNA sequencing, data processing and quality control were done as previously described for bone metastatic PC patients (*64*). Seurat was used to annotate cell subsets and calculate normalized gene score (*64*) of aerobic glycolysis (*65*) in tumor cells and M1-macrophage polarization status (AK3, APOL1, APOL2, APOL3, APOL6, ATF3, BCL2A1, BIRC3, CCL5, CCL15, CCL19, CCL20, CCR7, CHI3L2, CSPG2, CXCL9, CXCL10, CXCL11, ECGF1, EDN1, FAS, GADD45G, HESX1, HSD11B1, HSXIAPAF1, IGFBP4, IL6, IL12B, IL15, IL2RA, IL7R, IL15RA, INDO, INHBA, IRF1, IRF7, OAS2, OASL, PBEF1, PDGFA, PFKFB3, PFKP, PLA1A, PSMA2, PSMB9, PSME2, PTX3, SLC21A15, SLC2A6, SLC31A2, SLC7A5, SPHK1, TNF, TRAIL; (*67*)) of myeloid cells.

### *In vivo* macrophage depletion studies

To elucidate a role for macrophages in driving anti-tumor responses *in vivo*, Pb-Cre; PTEN^fl/fl^ Trp53^fl/fl^ mice with established prostate tumors were treated concurrently with clodronate (200 μg/mouse, *ip*, once weekly), which is known to deplete macrophages *in vivo,* and combinations of degarelix (0.625 mg/mouse, *sc*, single dose) + copanlisib (14 mg/kg, *iv*, every alternate day) + aPD-1 (200 μg/mouse, *ip*, every alternate day) for 28 days. Prostate tumor growth kinetics and immune profiling were done, as described in relevant subsections.

### *Ex vivo* cytokine array

*Ex vivo* CM were collected and analyzed using proteome profiler mouse XL cytokine array kit. Each blot was normalized using internal positive control and % change in cytokine secretion was calculated relative to untreated group.

### *In vivo* transcriptomic and pathway enrichment analysis

Pb-Cre; PTEN^fl/fl^ Trp53^fl/fl^ mice with established prostate tumors were treated with degarelix (0.625 mg/mouse, *sc*, single dose) + copanlisib (14 mg/kg, *iv*, every alternate day) + aPD-1 (200 μg/mouse, *ip*, every alternate day) combination for 28 days. Prostate tumors were harvested following treatment and RNA isolated using RNAeasy mini kits. Paired end sequencings were performed for 30 million reads on mRNA using NovaSeq 6000 platform. FASTQ raw files were generated by the sequencer, further pre-processed to discard the adaptor sequences using Cutadapt (*68*), and low-quality reads were eliminated. Then, filtered reads were mapped to mouse genome-mm9 using STAR aligner (*69*). STAR counts were used for differential expression analysis using DESeq2 R Bioconductor package (*70*). Pathway enrichment analysis was performed using Reactome tool on DEG datasets to discover mechanism for anti-tumor response and resistance to treatment (*71*).

### Data Analysis

Data was analyzed using GraphPad Prism 7 (GraphPad Software Inc.). Statistical analysis was performed using one-way analysis of variance (ANOVA) and Bonferroni post-test with p<0.05 level of significance. Bioinformatic data were analyzed using Wilcoxon ran-sum test. Chi-square test was performed to compare % ORR and % partial response.

## Acknowledgements

We thank Drs. Andrew Hsieh, Scott Oakes and Sam Grimaldo for providing helpful suggestions for this manuscript.

## Funding

This work was supported by;

- Prostate Cancer Foundation Challenge Award grant 16CHAL12 (AP)
- National Cancer Institute grant P30CA014599 (EM, MZ, XF)

## Author contributions

Conceptualization: KC, HMH, RK, AK, AP

Methodology: KC, HMH, RK, SM, RH, AK, EA, GAE, KH, ML, KB, CD, ASo, EM, MZ, XF

Investigation: KC, HMH, RK, SM, RH, AK

Visualization: KC, HMH, RK, SM, RH, AK, ASw, SS, DBS, AP

Funding acquisition: AP

Project administration: KC, HMH, AP Supervision: KC, AP

Writing – original draft: KC, HMH, RK, BL, TH, SM, JS, SR, DBS, AP

## Competing interest

Dr. Patnaik has received research funding from Bristol Myers Squibb.

## Data and materials availability

The authors declare that the data and materials supporting the findings of this study are available within the Article and its Supplementary Information.

## Supplementary figures

**Fig. S1.**
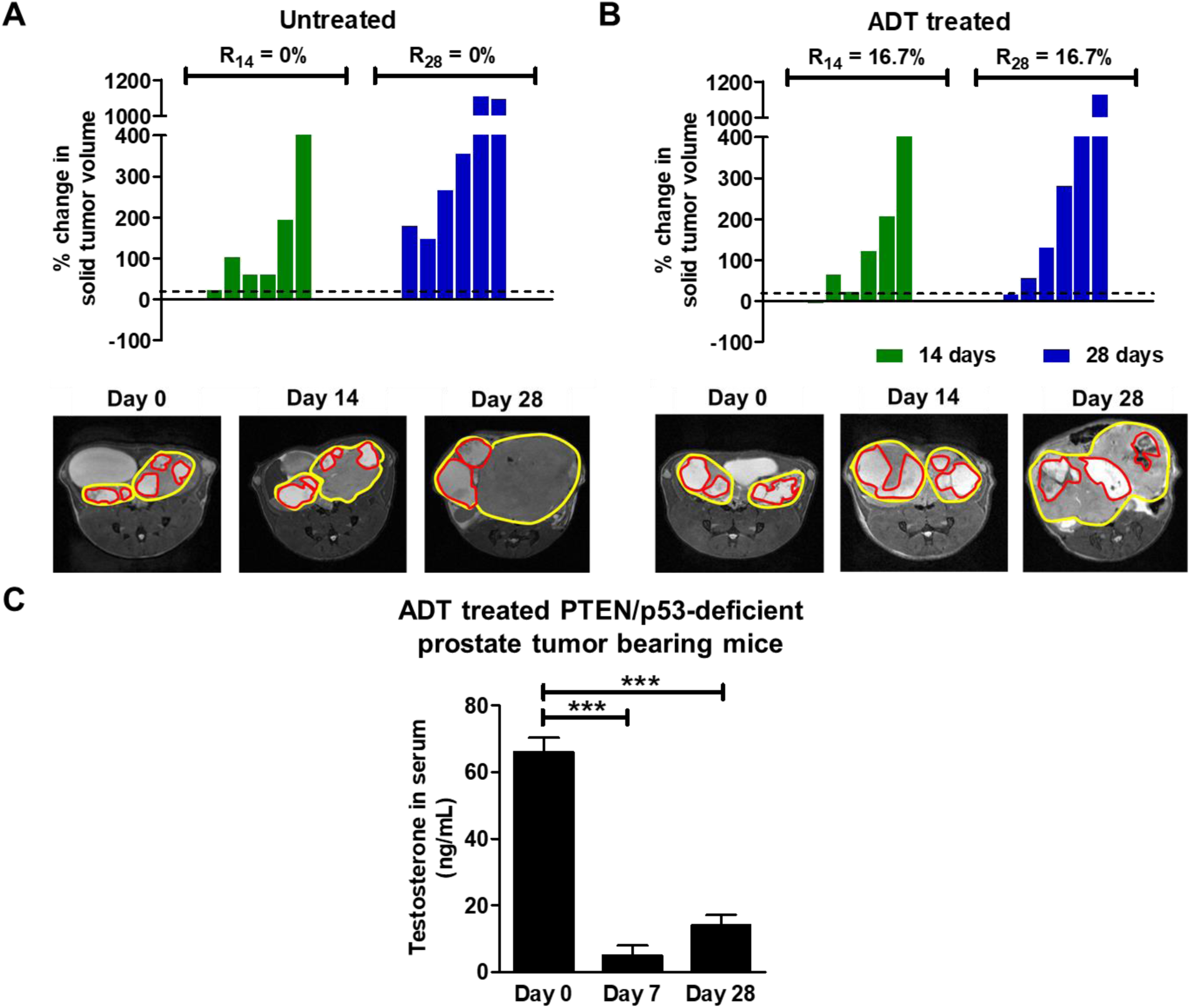
The majority of Pb-Cre; PTEN^fl/fl^ Trp53^fl/fl^ mice are *de novo* resistant to ADT. **(A-B)** Pb-Cre; PTEN^fl/fl^ Trp53^fl/fl^ mice with established prostate tumors were randomized to untreated or degarelix (ADT, 0.625 mg, single dose) for 4 weeks. Tumor volumes were non-invasively monitored by MRI, and % ORR at days 14 (R_14_) and 28 (R_28_) were determined as described in Methods. The % change in solid tumor volume is represented by waterfall plots for untreated **(A)** and ADT-treated groups **(B)**. **(C)** Sera were collected from mice at 7 and 28 days post-treatment, and analyzed for testosterone levels by ELISA. n=6-8 mice per group. Significances/p-values were calculated by Chi-square test (panel A and B, relative to untreated), one-way ANOVA (panel C) and indicates as follows, ***p<0.001.

**Fig. S2.**
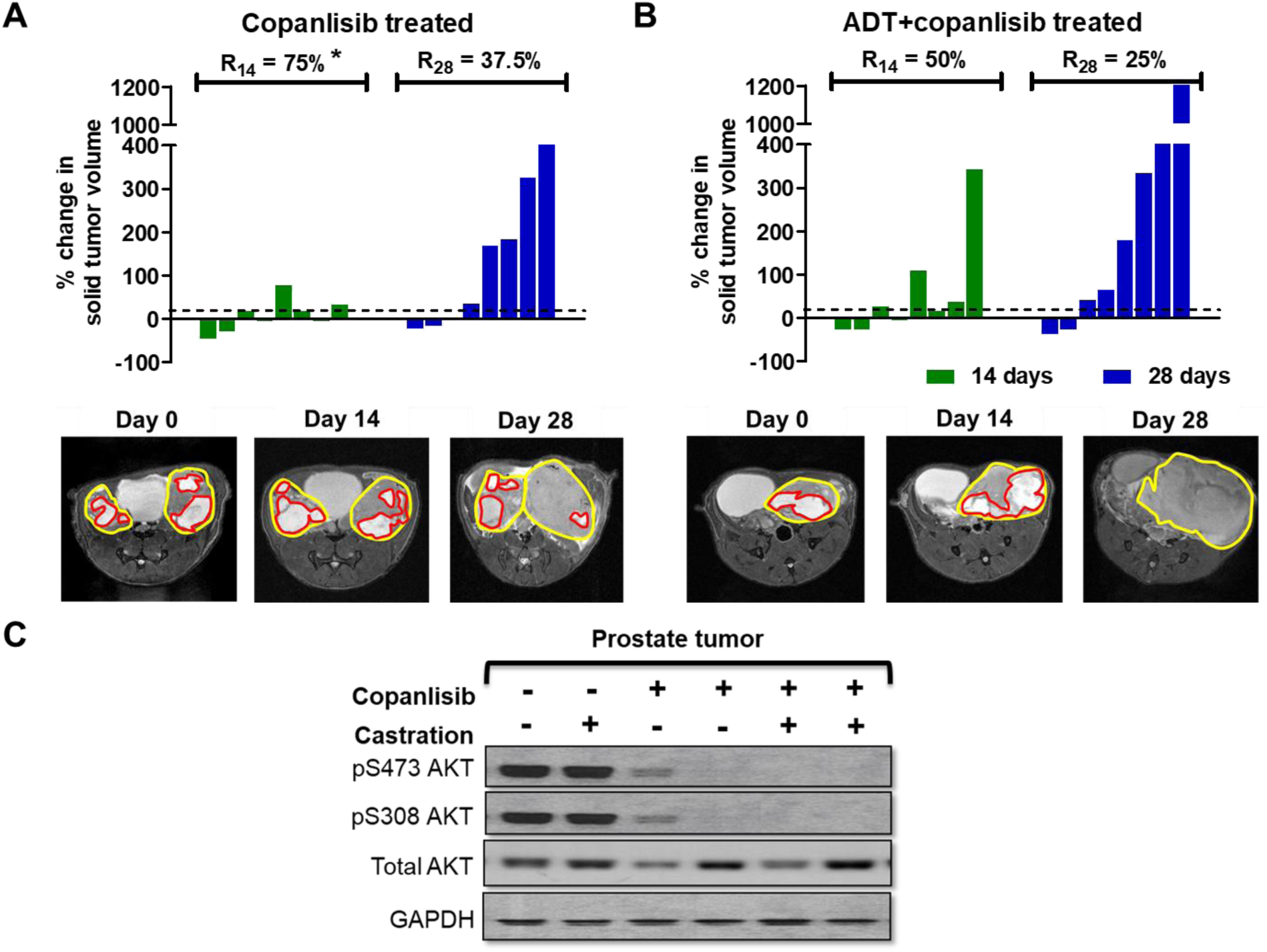
ADT/PI3Ki combination therapy halts prostate tumor growth up to 14 days, followed by development of resistance in majority of Pb-Cre; PTEN^fl/fl^ Trp53^fl/fl^ mice. **(A-B)** Pb-Cre; PTEN^fl/fl^ Trp53^fl/fl^ mice bearing established tumors were treated with copanlisib (14 mg/kg, *iv*, every alternate day), singly or in combination with ADT (degarelix, 0.625 mg, single dose). Tumor volumes were non-invasively monitored by MRI and % ORR at days 14 (R_14_) and 28 (R_28_) were determined as described in Methods. The % change in solid tumor volume is represented by waterfall plots for copanlisib **(A)** and ADT + copanlisib treated groups **(B)**. n=6-8 mice per group. Statistical analysis was performed using Chi-square test. *p<0.05 vs untreated. **(C)** Tumor extracts were harvested following 4 weeks of treatment with the indicated drug(s), and analyzed by western blotting for PI3K activation status. n=6-8 mice per group. Significances/p-values were calculated by Chi-square test (panel A and B, relative to untreated) and indicates as follows, *p<0.05.

**Fig. S3.**
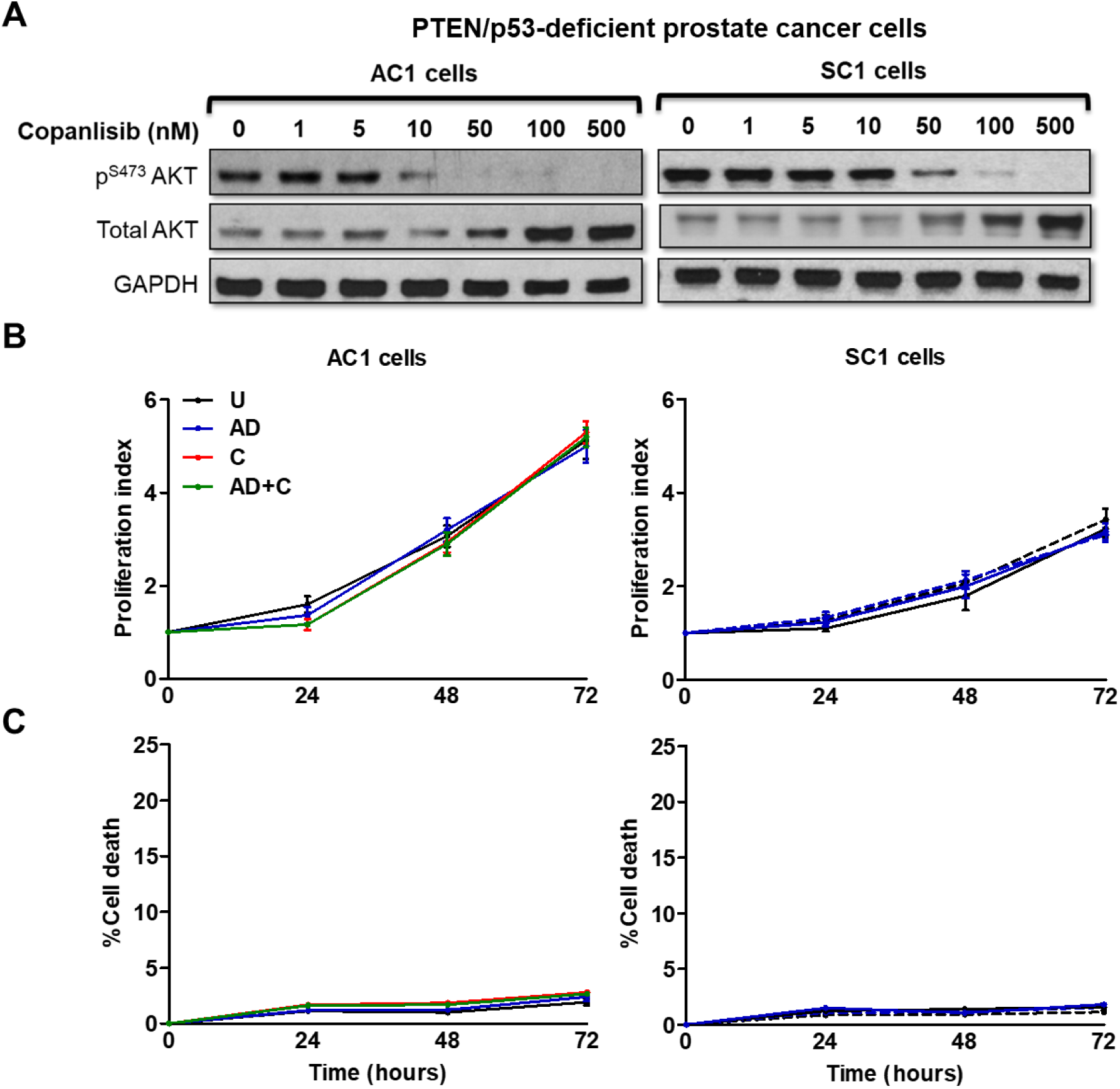
PI3Ki treatment with concurrent androgen depletion does not alter proliferation and survival of PTEN/p53-deficient murine PC cells *in vitro*. **(A)** PTEN/p53-deficient GEMM tumor-derived PC cell lines (AC1/SC1) were treated with copanlisib (C, 0-500 nM, 24 hours) in a dose-dependent manner and western blot analyses were performed for indicated proteins to determine IC_90_. **(B-C)** AC1/SC1 cells were treated with copanlisib (IC_90_=100 nM, for 24, 48 and 72 hours) in a time-dependent manner in normal and AD conditions. **(B)** Proliferation index was calculated by counting cell number at the end of treatment, relative to their baseline seeding density. **(C)** Cells were stained with annexin V antibody/propidium iodide and % cell death was determined via flow cytometry. U=untreated group. n=3 independent experiments. Significances/p-values were calculated by one-way ANOVA.

**Fig. S4.**
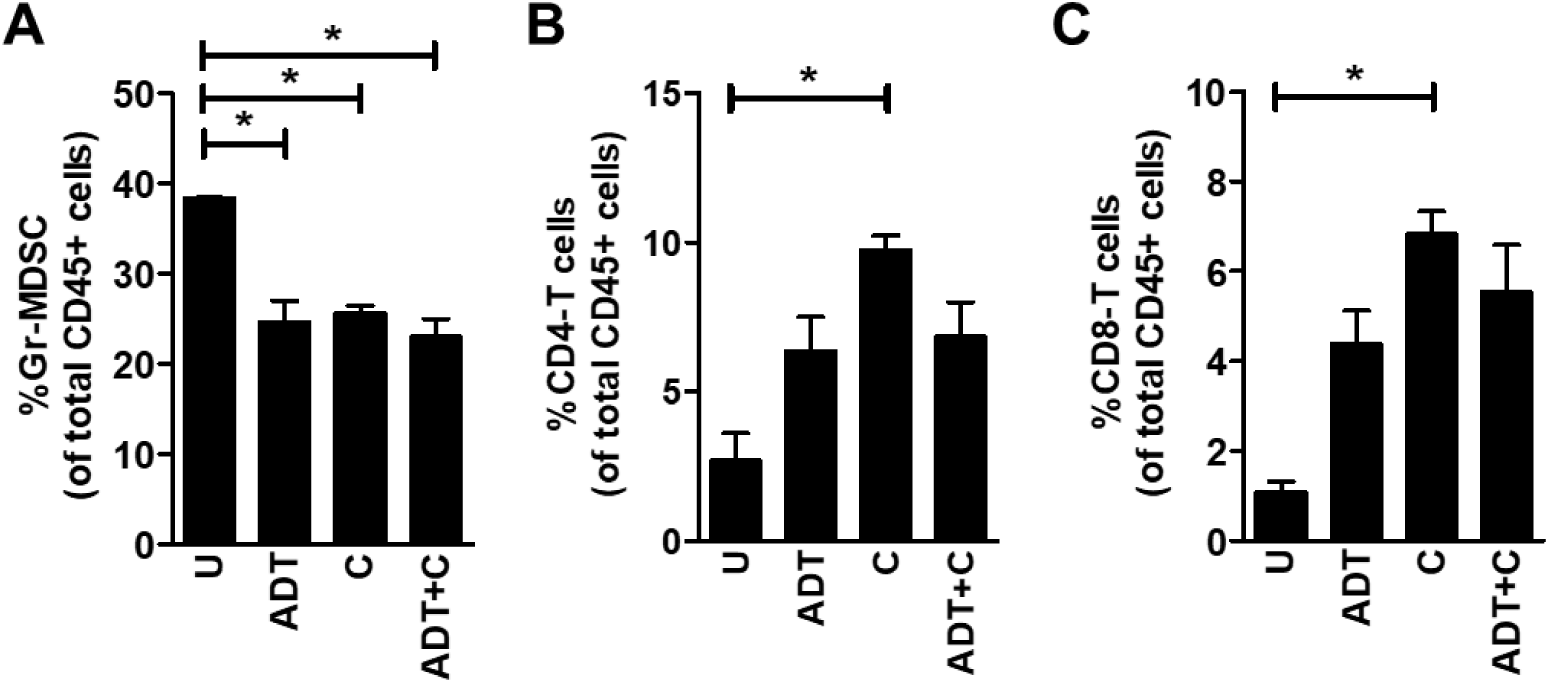
ADT or PI3K inhibitor, singly or in combination, alleviated immunosuppression within the TME of PTEN/p53-deficient murine PC. **(A-C)** Pb-Cre; PTEN^fl/fl^ Trp53^fl/fl^ mice with established tumors were treated with degarelix (0.625 mg, single dose, ADT), copanlisib (14 mg/kg, *iv*, every alternate day, C) or their combination for 7 days. Tumor cell suspensions were collected and analyzed using flow cytometry. Bar graph demonstrates % frequency of granulocytic-myeloid derived suppressive cells (Gr-MDSC, CD45^+^CD11b^+^F4/80^-^Ly6g^+^ cells, **(A)**), % frequency of CD4-T cells (CD45^+^CD3^+^CD4^+^ cells, **(B)**) and % frequency of CD8-T cells (CD45^+^CD3^+^CD8^+^ cells, **(C)**). U=untreated group. n=2 mice per group. Significances/p-values were calculated by one-way ANOVA and indicates as follows, *p<0.05.

**Fig. S5.**
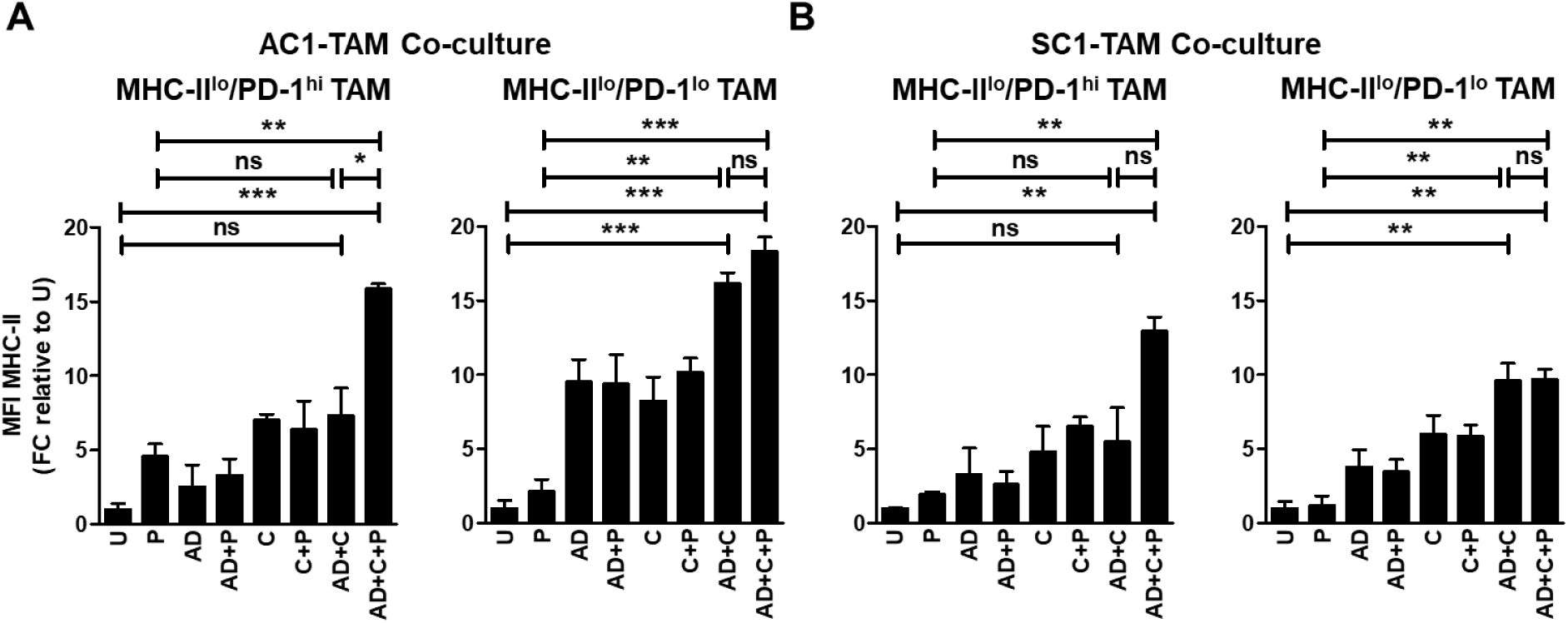
*Ex vivo* AD + PI3Ki + PD-1 antibody treatment activates MHC-II^lo^ TAM when co-cultured with PTEN/p53-deficient murine prostate tumor cells. **(A-B)** TAM subsets (MHC-II^lo^/PD-1^hi/lo^) were sorted from PTEN/p53-deficient prostate tumors using flow cytometry, and co-cultured with PTEN/p53-deficient prostate tumor cells under normal and AD conditions. These co-cultures were treated with copanlisib (C, 100 nM), PD-1 antibody (P, 10 μg/mL) or their combination for 24 hours. Bar graphs demonstrate MHC-II expression on MHC-II^lo^/PD-1^hi/lo^ TAM following co-culture with AC1 **(A)** and SC1 **(B)** cells in the presence of indicated treatments, relative to untreated (U). FC=fold change. n=2 independent experiments. Significances/p-values were calculated by one-way ANOVA and indicates as follows, *p<0.05, **p<0.01 and ***p<0.001; ns = not statistically significant.

**Fig. S6.**
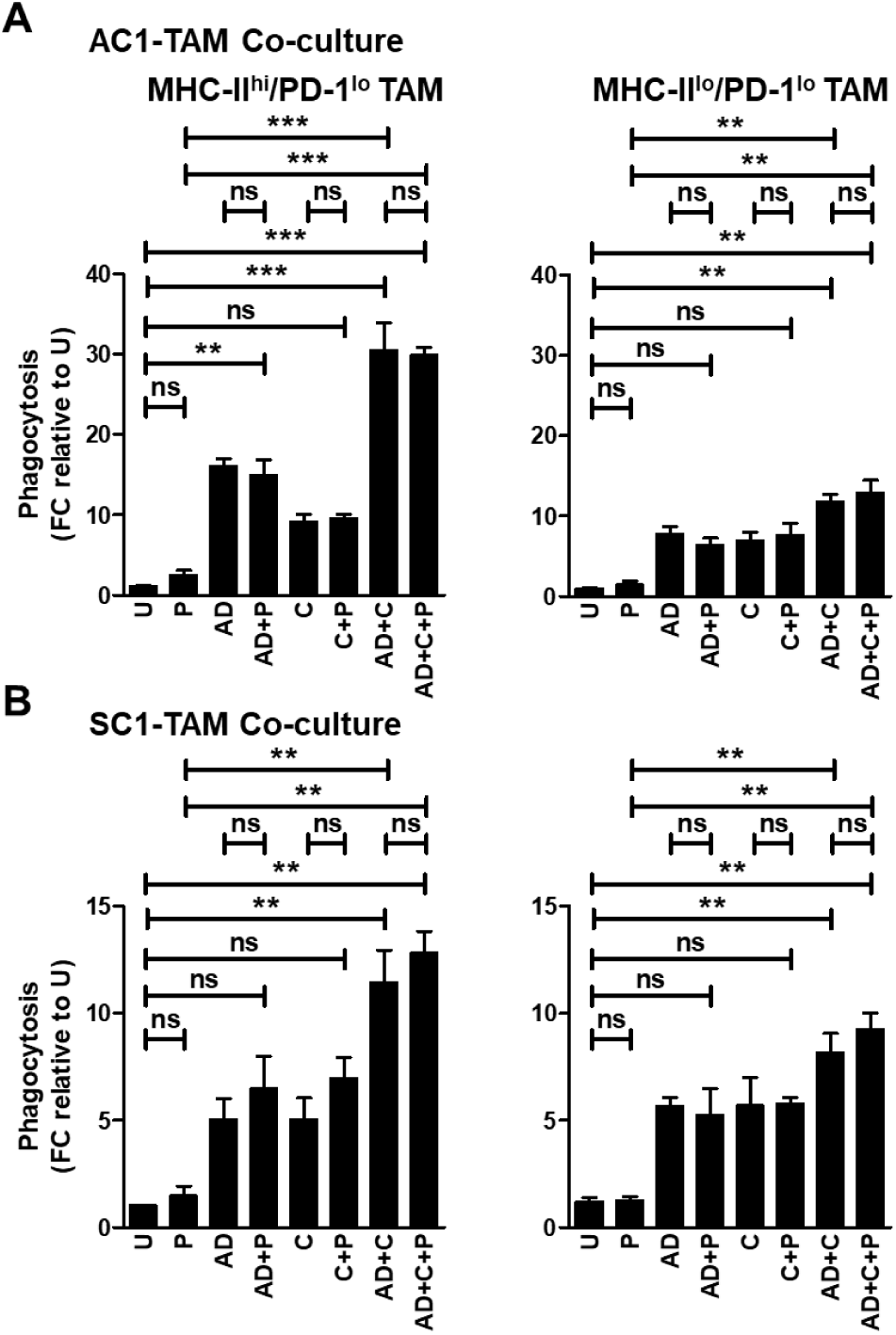
The addition of PD-1 blockade to androgen depletion/PI3Ki therapy does not alter phagocytic capacity of PD-1^lo^ macrophages. **(A)** TAM subsets (MHC-II^hi/lo^/PD-1^lo^) sorted from PTEN/p53-deficient prostate tumors using flow cytometry were co-cultured with CTV dye stained-AC1/SC1 cells under normal and AD conditions. These co-cultures were treated with copanlisib (C, 100 nM), PD-1 antibody (P, 10 μg/mL) or their combination for 24 hours. Bar graphs demonstrate fold change (FC) in phagocytic activity of MHC-II^hi/lo^/PD-1^lo^ expressing TAM relative to untreated group (U) in AC1 **(B)** and SC1 **(C)** cells. n=2 independent experiments. Significances/p-values were calculated by one-way ANOVA and indicates as follows, **p<0.01 and ***p<0.001; ns = not statistically significant.

**Fig. S7.**
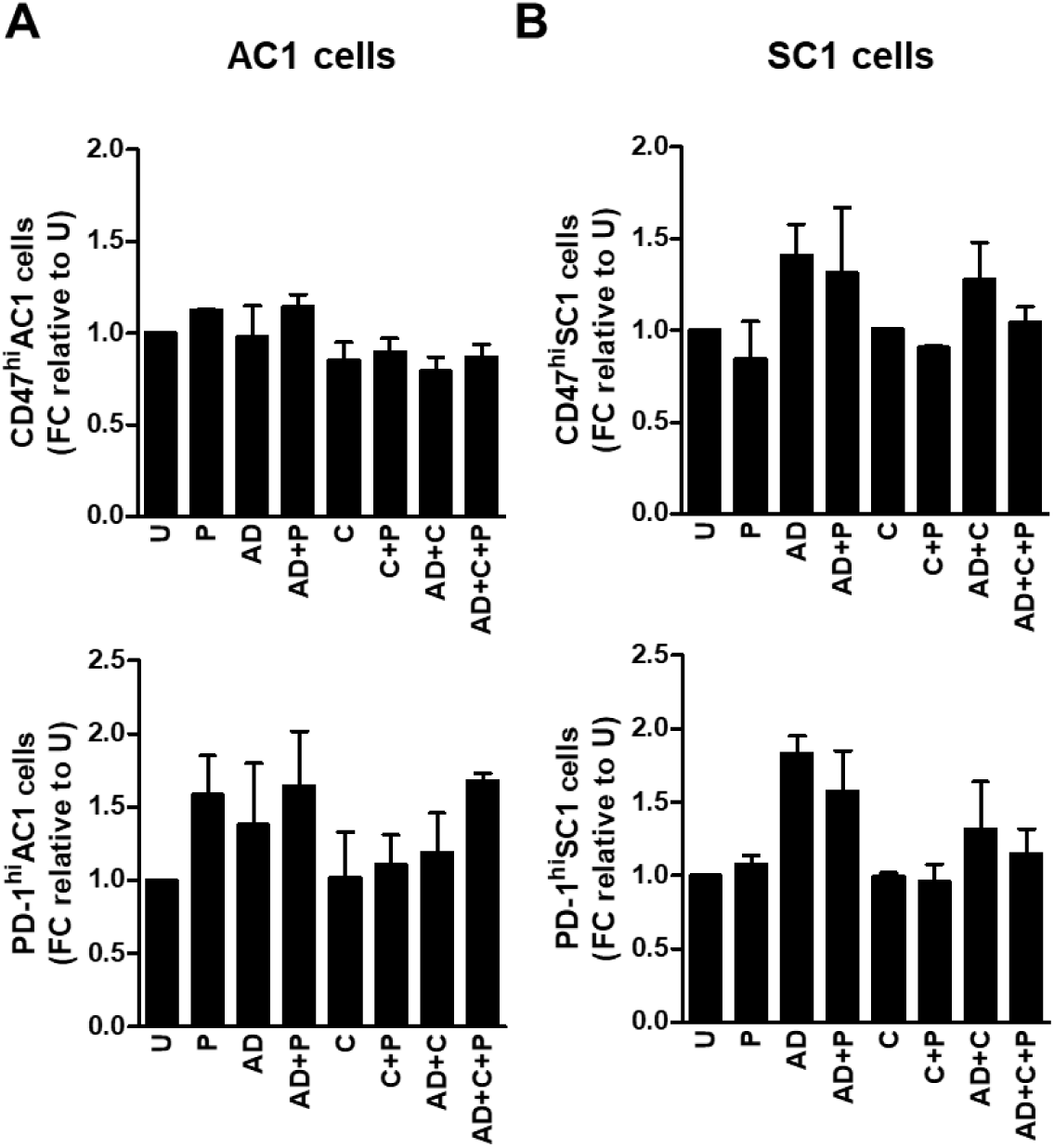
The combination of androgen depletion, PI3Ki and aPD-1 blockade does not alter phagocytic checkpoint expression on PTEN/p53-deficient prostate tumor cells. **(A-B)** AC1/SC1 cells were treated with copanlisib (C, 100 nM), PD-1 antibody (P, 10 μg/mL) or their combination under normal and AD conditions for 24 hours. Bar graphs demonstrate phagocytosis inhibitory checkpoints (CD47 and PD-1) expression on AC1 **(A)** and SC1 **(B)** cells following indicated treatments. U=untreated group. FC=fold change. n=2 independent experiments. Significances/p-values were calculated by one-way ANOVA.

**Fig. S8.**
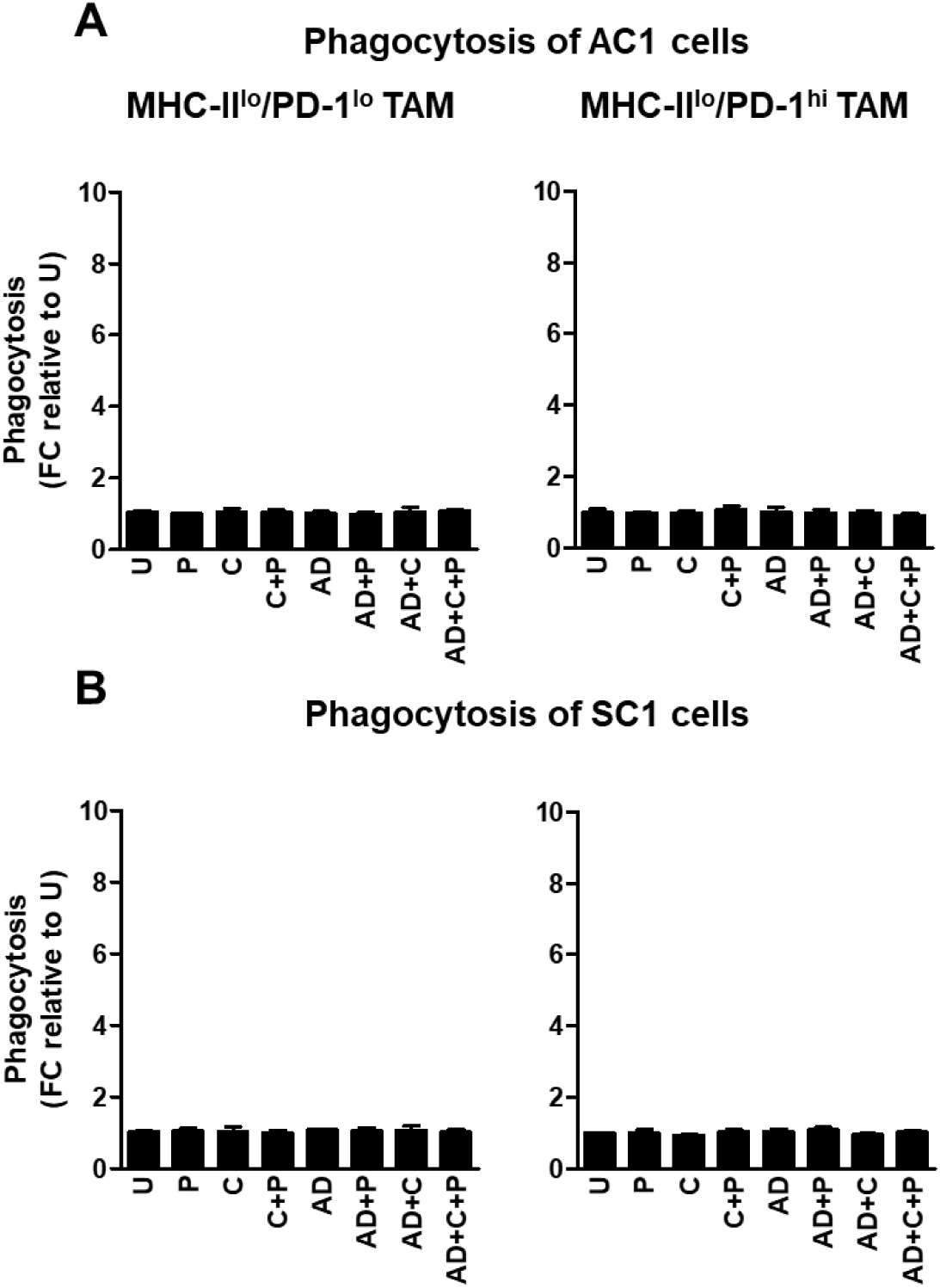
Androgen depletion, singly and in combination with aPD-1, did not alter phagocytosis activity of inactivated MHC-II^lo^/PD-1^lo^ and MHC-II^lo^/PD-1^hi^ TAM subsets. **(A-B)** Pre-treatment of FACS-sorted TAM subsets from PTEN/p53-deficient prostate tumors with copanlisib (C, 100 nM), PD-1 antibody (P, 10 μg/mL) or their combinations under normal and AD conditions for 24 hours. After PBS wash, treated TAM were co-cultured with CTV dye stained-AC1/SC1 cells for 2 hours. Bar graphs demonstrate fold change (FC) in phagocytosis of AC1 **(A)** and SC1 **(B)** cells by MHC-II^lo^/PD-1^hi/lo^ expressing TAM, relative to untreated group (U). n=2 independent experiments. Significances/p-values were calculated by one-way ANOVA.

**Fig. S9.**
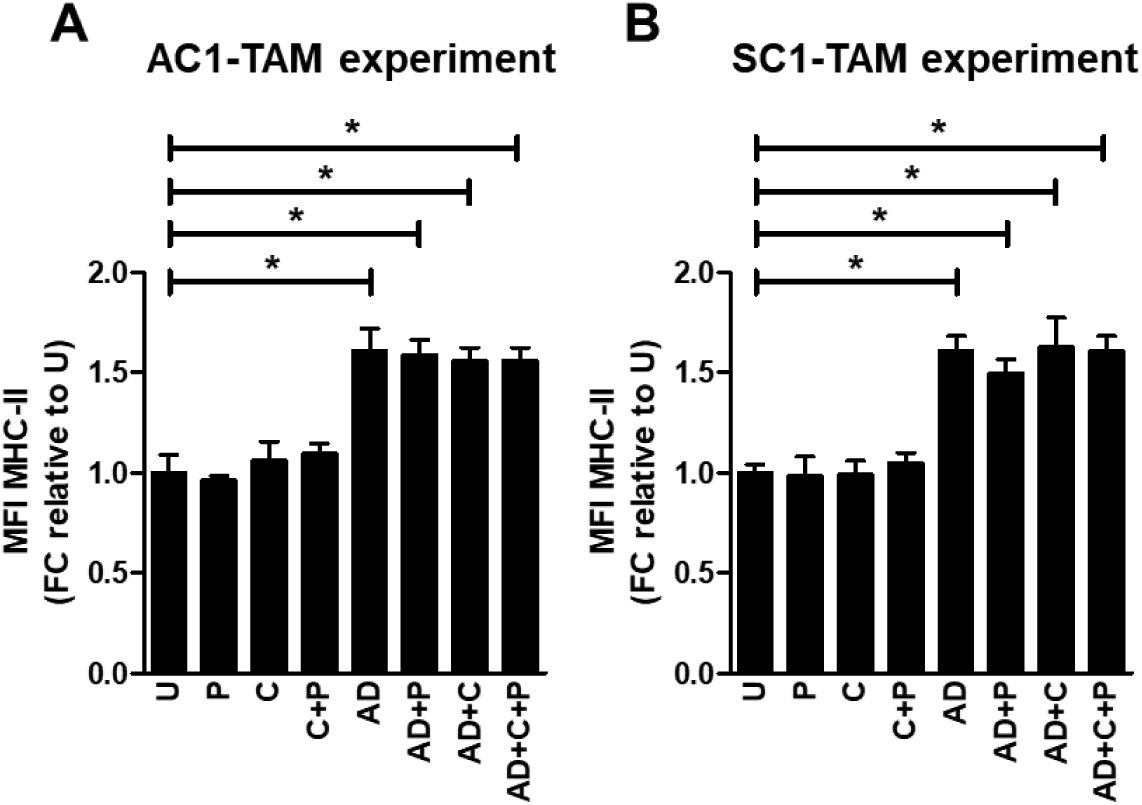
Androgen depletion, not PI3Ki or aPD1, directly enhances TAM activation within the TME of PTEN/p53-deficient PC. **(A-B)** FACS-sorted TAM from PTEN/p53-deficient prostate tumors were directly treated with copanlisib (C, 100 nM), PD-1 antibody (P, 10 μg/mL) or their combinations under normal and AD conditions for 24 hours. Bar graphs demonstrate MHC-II expression on TAM following co-culture with AC1 **(A)** and SC1 **(B)** cells for 2 hours, in the presence of indicated treatments. U=untreated group. FC=fold change. n=2 independent experiments. Significances/p-values were calculated by one-way ANOVA and indicates as follows, *p<0.05.

**Fig. S10.**
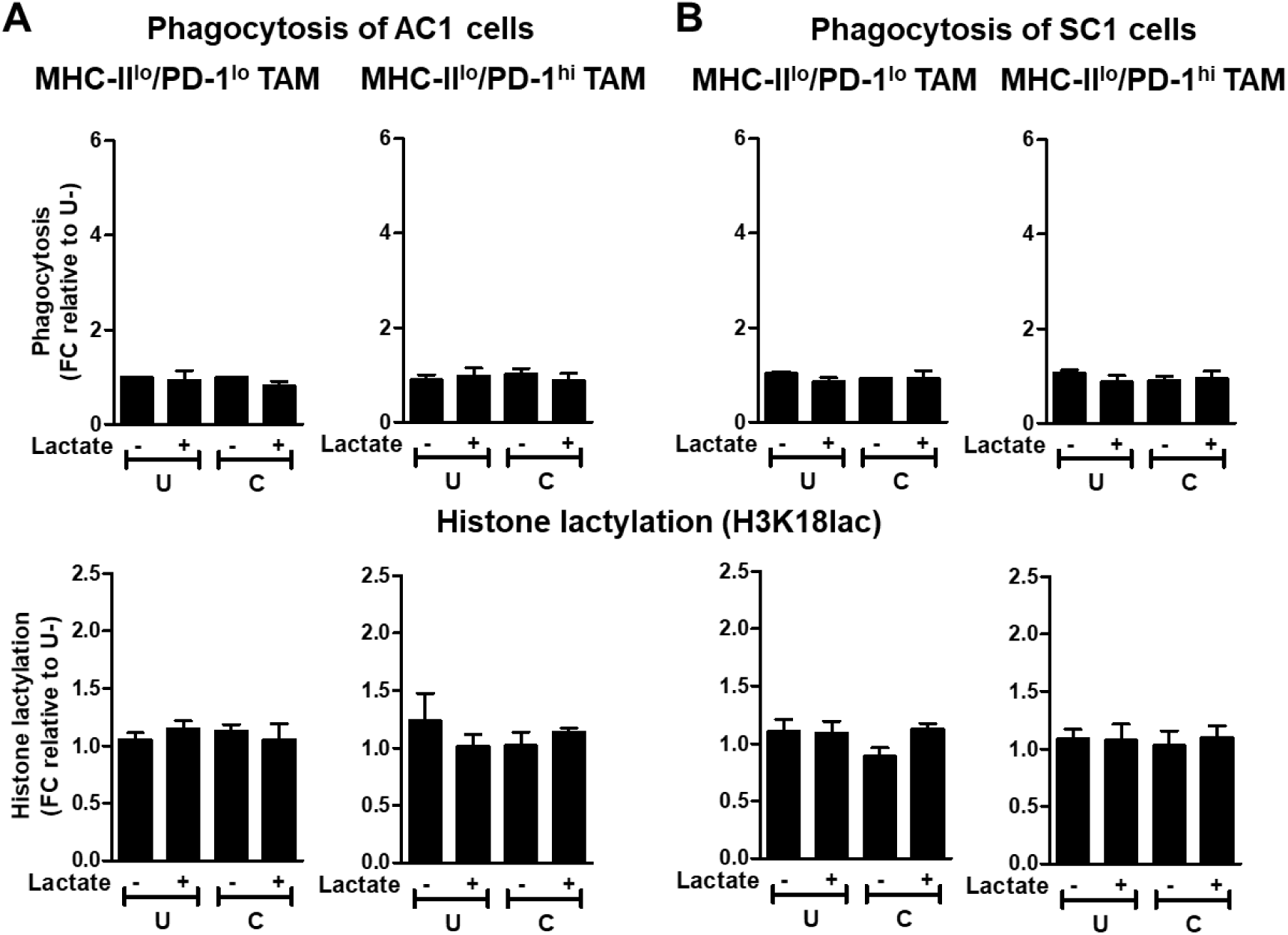
PI3Ki does not alter phagocytosis/histone lactylation status of MHC-II^lo^/PD-1^lo^ TAM and MHC-II^lo^/PD-1^hi^ TAM. **(A-B)** Single cell suspension of PTEN/p53-deficient prostate GEMM tumors were treated with copanlisib (C, 100 nM) and CM was collected 24 hours following treatment. FACS-sorted TAM from PTEN/p53-deficient prostate tumors were cultured *ex vivo* with CM for 24 hours in presence or absence of lactate (100 nmol/μL). After PBS wash, TAM were co-cultured with CTV dye stained-AC1/SC1 cells for 2 hours. Bar graphs demonstrate fold change (FC) in phagocytic activity and histone lactylation status of inactivated MHC-II^lo^/PD-1^lo^ and MHC-II^lo^/PD-1^hi^ TAM following incubation with AC1 **(A)** and SC1 **(B)** cells, relative to untreated group. n=2 independent experiments. Significances/p-values were calculated by one-way ANOVA.

**Fig. S11.**
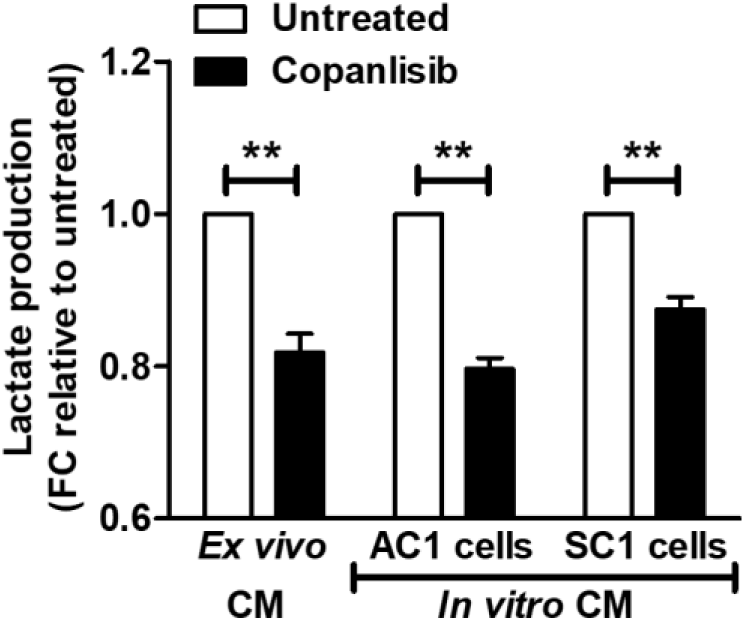
PI3Ki inhibits lactate secretion from PTEN/p53-deficient prostate tumor cells within TME. Single cell suspensions of PTEN/p53-deficient prostate GEMM tumors or tumor-derived AC1/SC1 cells were treated with copanlisib (100 nM) for 24 hours. *Ex vivo* or *in vitro* CM were collected from these groups and analyzed for lactate content using colorimetry kits. n=3 independent experiments. Significances/p-values were calculated by one-way ANOVA and indicates as follows, **p<0.01.

**Fig. S12.**
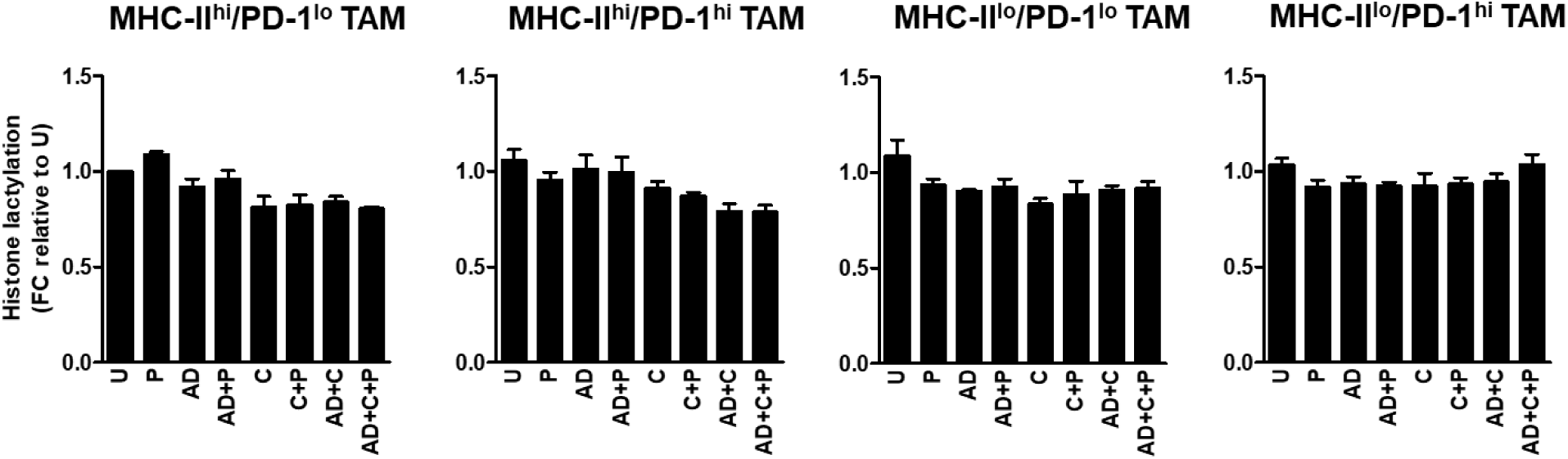
Direct *ex vivo* treatment of TAM with PI3Ki, singly and in combination with PD-1 antibody and/or androgen depletion does not alter their histone lactylation profile. FACS-sorted TAM from PTEN/p53-deficient prostate tumors were directly treated with copanlisib (C, 100 nM), PD-1 antibody (P, 10 μg/mL) or their combinations under normal and AD conditions for 24 hours. Bar graphs demonstrate histone lactylation status of MHC-II^hi/lo^/PD-1^hi/lo^ expressing TAM in response to indicated treatments. U=untreated group. FC=fold change. n=2 independent experiments. Significances/p-values were calculated by one-way ANOVA.

**Fig. S13.**
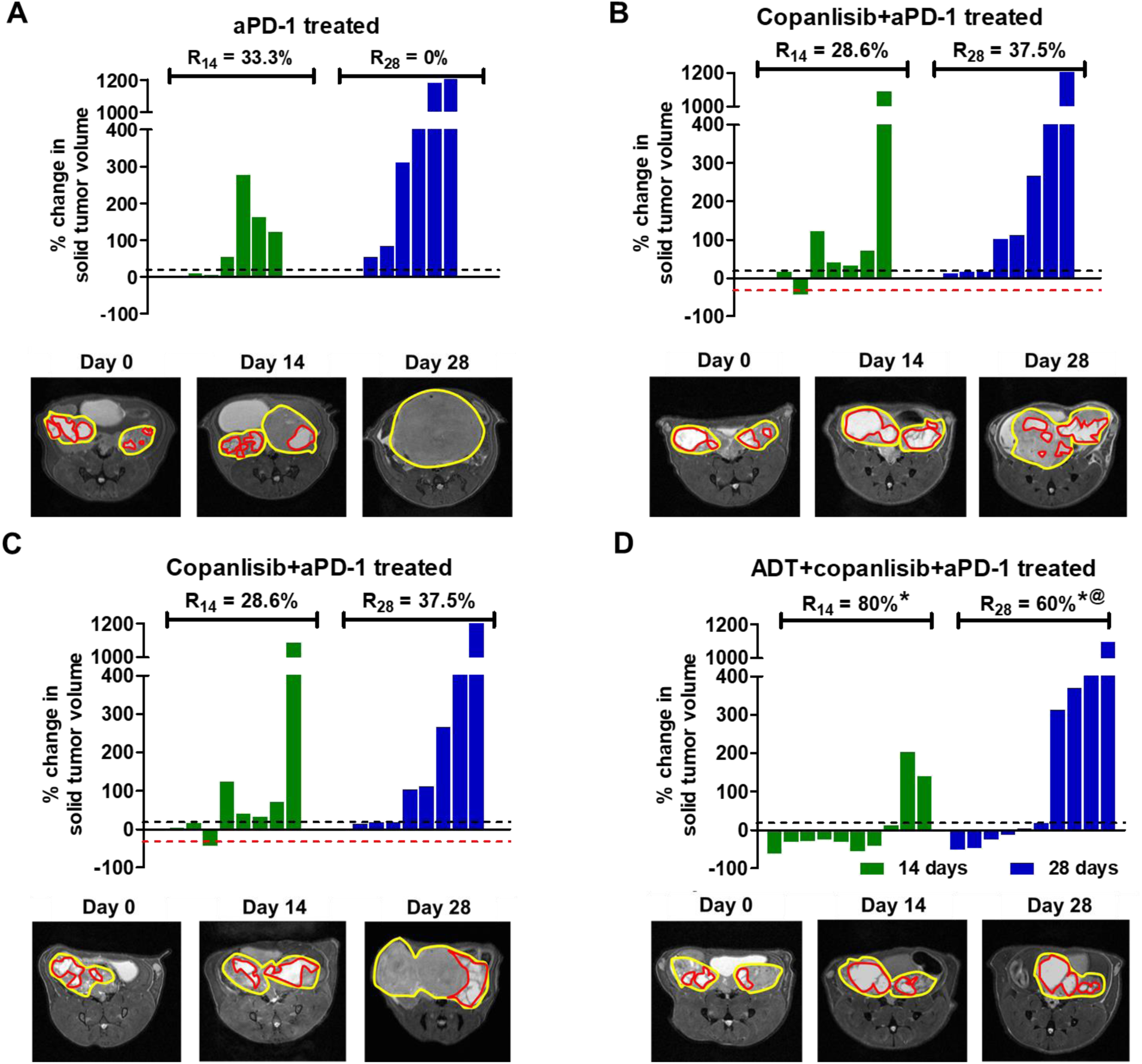
ADT + PI3Ki + aPD-1 induces tumor control up to 14 days followed by development of resistance in 40% of Cre-Pb; PTEN^fl/fl^ TP53^fl/fl^ mice. **(A-D)** Pb-Cre; PTEN^fl/fl^ Trp53^fl/fl^ mice with established tumors were treated with PD-1 antibody (aPD-1, 100 μg/mouse, *ip*, every alternate day) alone or in combination with ADT (degarelix, 0.625mg, single dose) or copanlisib (14 mg/kg, *iv*, every alternate day) or ADT + copanlisib. Tumor volumes were non-invasively monitored by MRI and % ORR at days 14 (R_14_) and 28 (R_28_) were determined, as described in Methods. The % change in solid tumor volume are represented by waterfall plot for the following treated groups: aPD-1 **(A)**, ADT + aPD-1 **(B)**, copanlisib + aPD-1 **(C)** and ADT + copanlisib + aPD-1 **(D)**. n=6-10 mice per group. Significances/p-values were calculated by Chi-square test and indicated as follows, *p<0.05 and ^@^p<0.05 (relative to untreated and aPD-1 treated groups, respectively).

**Fig. S14.**
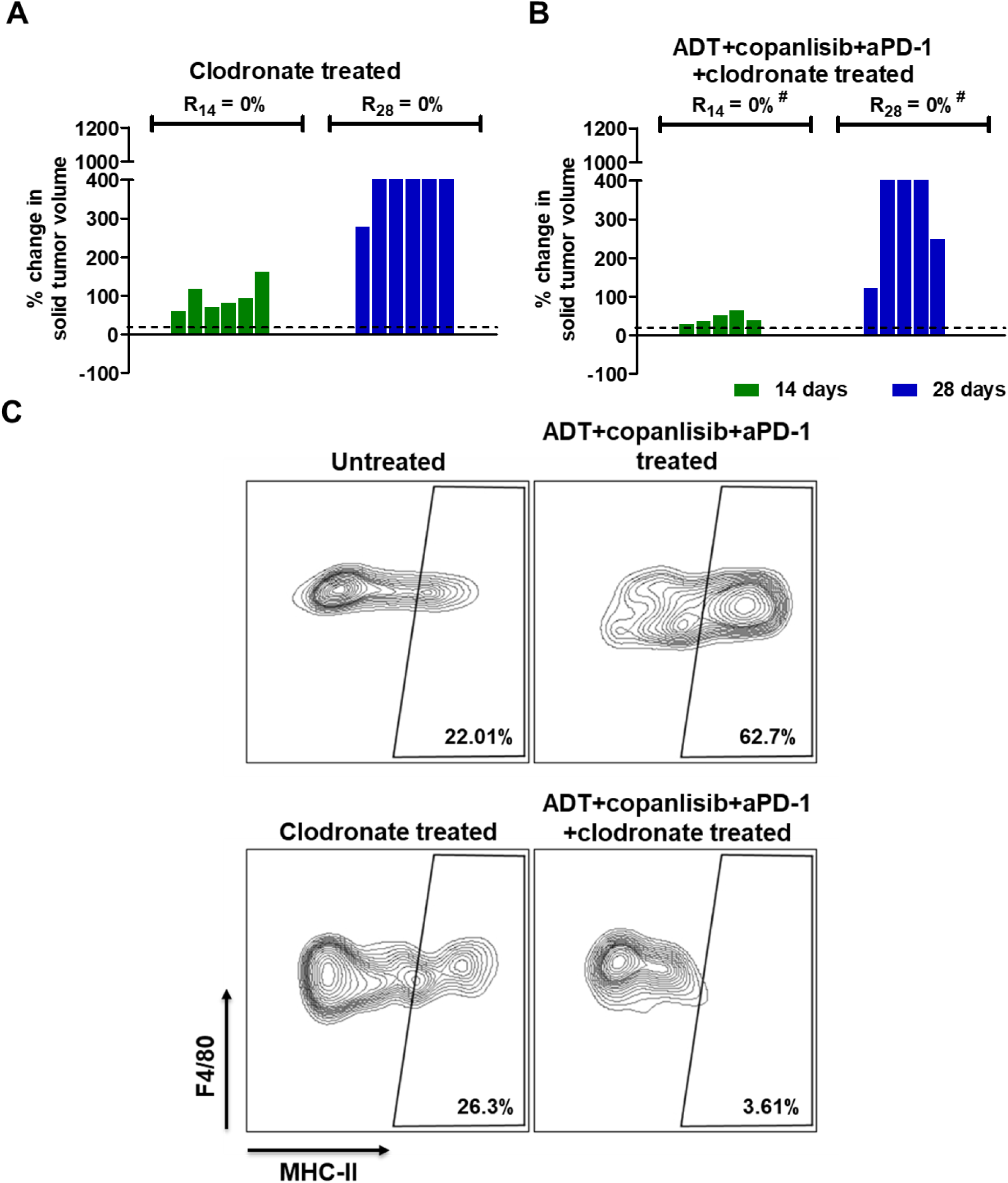
Depletion of activated TAM abrogates anti-cancer response elicited by ADT + PI3Ki + PD-1 antibody treatment in the PTEN/p53-deficient murine prostate GEMM tumors. **(A-B)** Pb-Cre; PTEN^fl/fl^ Trp53^fl/fl^ mice with established prostate tumors were treated with clodronate (which depletes activated macrophages, 200 μg/mouse, *ip*, every week) alone or in combination with ADT (degarelix, 0.625mg, single dose) + copanlisib (14 mg/kg, *iv*, every alternate day) + PD-1 antibody (aPD-1, 100 μg/mouse, *ip*, every alternate day). Tumor volumes were non-invasively monitored by MRI and % ORR at days 14 (R_14_) and 28 (R_28_) were determined as described in Methods. The % change in solid tumor volume are represented by waterfall plot for clodronate **(A)** and ADT + copanlisib + aPD-1 + clodronate **(B)** treated groups. **(C)** Flow cytometry plot represents frequency of MHC-II expressing TAM within GEMM tumors from mice treated with the indicated drugs, relative to untreated mice. n=5-6 mice per group. Significances/p-values were calculated by Chi-square test and indicates as follows, #p<0.05 (relative to ADT/copanlisib/aPD-1 combination treated group).

**Fig. S15.**
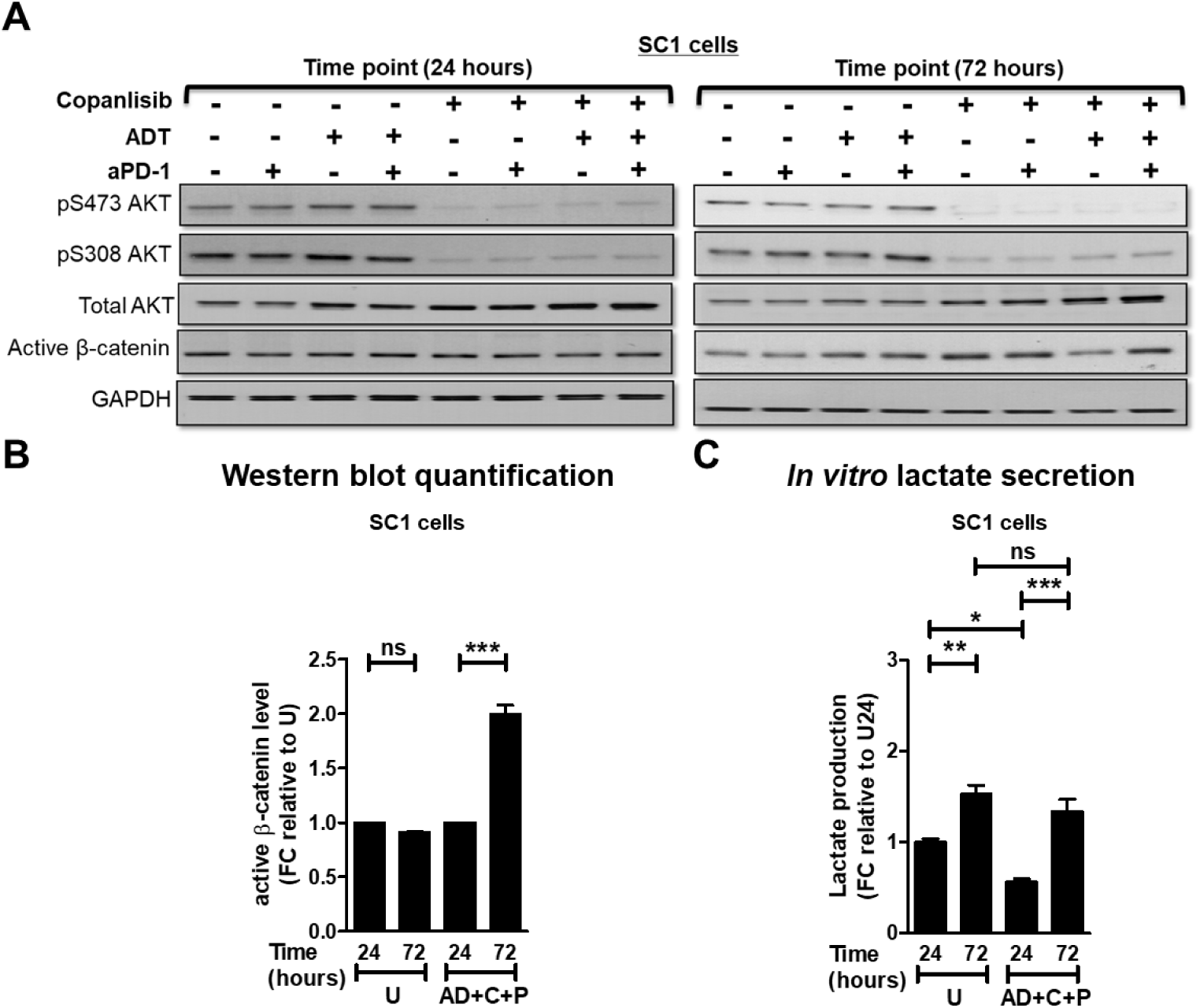
Long-term treatment of ADT + PI3Ki + aPD-1 activates Wnt/β-catenin pathway in murine PTEN/p53-deficient GEMM-derived SC1 cells. **(A-C)** SC1 cells were treated with copanlisib (C, 100 nM) + PD-1 antibody (P, 10 μg/mL) under AD condition for 24 and 72 hours. Western blot analyses **(A)** were performed for indicated protein markers on SC1 cell lysates, quantified using Image J software **(B)**, and lactate levels **(C)** were determined by fluorimetry in the indicated SC1 supernatant groups. U=untreated group. n=3 independent experiments. Significances/p-values were calculated by one-way ANOVA and indicates as follows, *p<0.05, **p<0.01 and ***p<0.001; ns = not statistically significant.

**Fig. S16.**
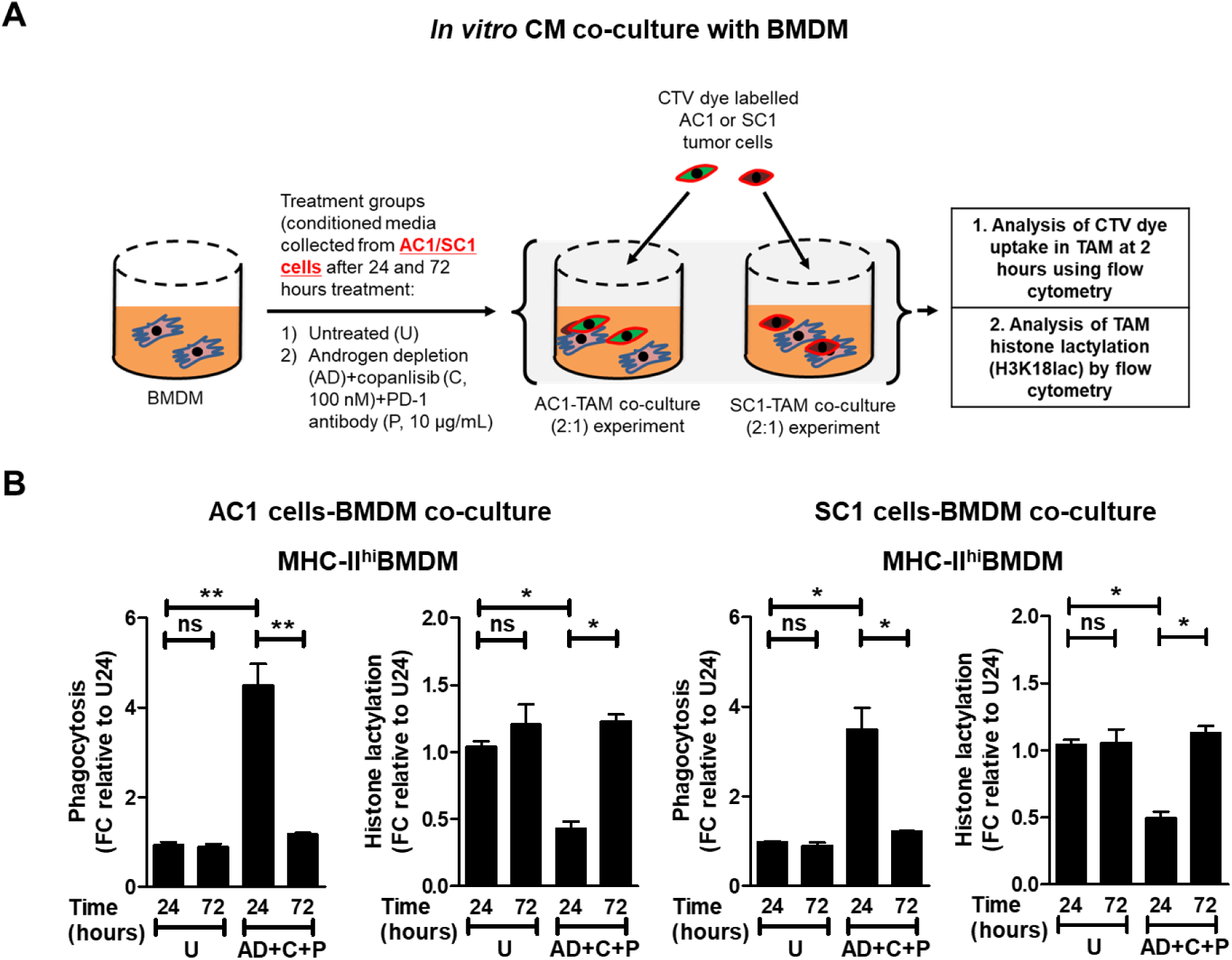
Feedback Wnt/β-catenin-pathway activation within murine PTEN/p53-deficient GEMM-derived PC cells following long-term ADT + copanlisib + aPD-1 treatment suppresses phagocytosis via increased histone lactylation within bone marrow derived macrophages (BMDM). **(A)** Schema illustrating phagocytosis experiment on *in vitro* CM. AC1/SC1 cells were treated with copanlisib (C, 100 nM) + PD-1 antibody (P, 10 μg/mL) under AD condition for 24 and 72 hours. *In vitro* CM were collected from these groups and co-cultured with BMDM for 24 hours. After PBS wash, these BMDM were co-cultured with CTV dye stained-AC1/SC1 cells for 2 hours. **(B)** Bar graphs demonstrate fold change (FC) in phagocytic activity and histone lactylation status of activated (MHC-II^hi^) BMDM when co-cultured with AC1/SC1 cells, relative to untreated group (U). n=3 independent experiments. Significances/p-values were calculated by one-way ANOVA and indicates as follows, *p<0.05 and **p<0.01; ns = not statistically significant.

## Supplementary table

**Table S1.**
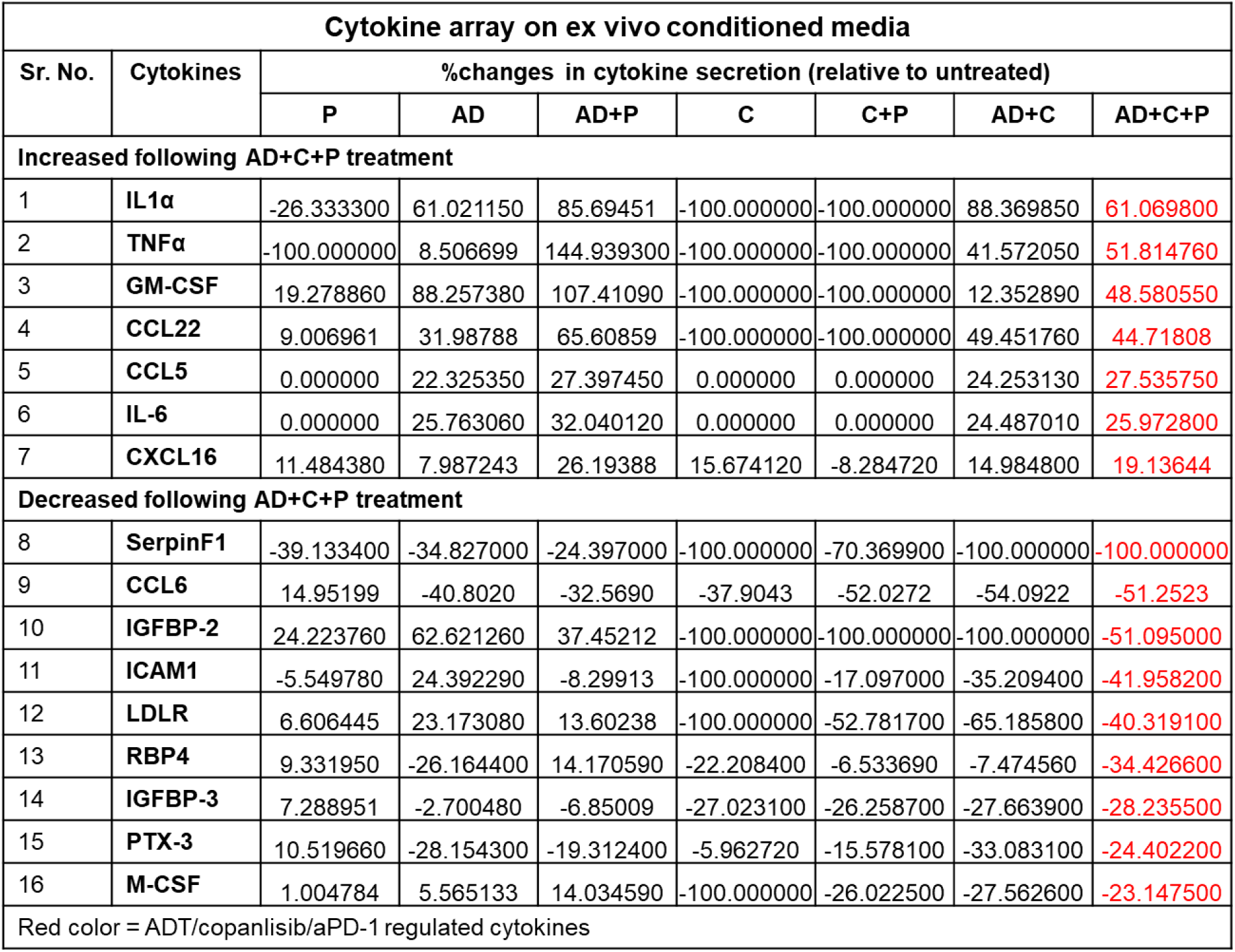
ADT + PI3Ki + PD-1 antibody leads to TAM activation within TME of PTEN/p53- deficient prostate tumors. Single cell suspension of PTEN/p53-deficient prostate tumors were treated with copanlisib (C, 100 nM), PD-1 antibody (P, 10 μg/mL) or their combination under normal and AD condition for 24 hours. *Ex vivo* conditioned media were collected from these groups and analyzed using cytokine array kits.

| Cytokine array on ex vivo conditioned media |  |  |  |  |  |  |  |  |
| --- | --- | --- | --- | --- | --- | --- | --- | --- |
| Sr. No. | Cytokines | %changes in cytokine secretion (relative to untreated) |  |  |  |  |  |  |
|  |  | P | AD | AD+P | C | C+P | AD+C | AD+C+P |
| Increased following AD+C+P treatment |  |  |  |  |  |  |  |  |
| 1 | IL1α | -26.333300 | 61.021150 | 85.69451 | -100.000000 | -100.000000 | 88.369850 | 61.069800 |
| 2 | TNFα | -100.000000 | 8.506699 | 144.939300 | -100.000000 | -100.000000 | 41.572050 | 51.814760 |
| 3 | GM-CSF | 19.278860 | 88.257380 | 107.41090 | -100.000000 | -100.000000 | 12.352890 | 48.580550 |
| 4 | CCL22 | 9.006961 | 31.98788 | 65.60859 | -100.000000 | -100.000000 | 49.451760 | 44.71808 |
| 5 | CCL5 | 0.000000 | 22.325350 | 27.397450 | 0.000000 | 0.000000 | 24.253130 | 27.535750 |
| 6 | IL-6 | 0.000000 | 25.763060 | 32.040120 | 0.000000 | 0.000000 | 24.487010 | 25.972800 |
| 7 | CXCL16 | 11.484380 | 7.987243 | 26.19388 | 15.674120 | -8.284720 | 14.984800 | 19.13644 |
| Decreased following AD+C+P treatment |  |  |  |  |  |  |  |  |
| 8 | SerpinF1 | -39.133400 | -34.827000 | -24.397000 | -100.000000 | -70.369900 | -100.000000 | -100.000000 |
| 9 | CCL6 | 14.95199 | -40.8020 | -32.5690 | -37.9043 | -52.0272 | -54.0922 | -51.2523 |
| 10 | IGFBP-2 | 24.223760 | 62.621260 | 37.45212 | -100.000000 | -100.000000 | -100.000000 | -51.095000 |
| 11 | ICAM1 | -5.549780 | 24.392290 | -8.29913 | -100.000000 | -17.097000 | -35.209400 | -41.958200 |
| 12 | LDLR | 6.606445 | 23.173080 | 13.60238 | -100.000000 | -52.781700 | -65.185800 | -40.319100 |
| 13 | RBP4 | 9.331950 | -26.164400 | 14.170590 | -22.208400 | -6.533690 | -7.474560 | -34.426600 |
| 14 | IGFBP-3 | 7.288951 | -2.700480 | -6.85009 | -27.023100 | -26.258700 | -27.663900 | -28.235500 |
| 15 | PTX-3 | 10.519660 | -28.154300 | -19.312400 | -5.962720 | -15.578100 | -33.083100 | -24.402200 |
| 16 | M-CSF | 1.004784 | 5.565133 | 14.034590 | -100.000000 | -26.022500 | -27.562600 | -23.147500 |

Red color = ADT/copanlisib/aPD-1 regulated cytokines

**Table S2.** Chemical and reagents.

| Sr. No. | Chemicals and reagents | Catalogue no. | Vendor/provider |
| --- | --- | --- | --- |
| 1 | Deagerlix | HY16168A | MedChemExpress |
| 2 | PD-1 antibody | mPD-1-D265A | Bristol Myers Squibb |
| 3 | Copanlisib | BAY 80-6946 | Selleckchem |
| 4 | Serum testosterone determination kits | ADI-900-176 | Enzo Life Sciences |
| 5 | PrEGM media | CC-3166 | Lonza |
| 6 | Fetal bovine serum (FBS) | 100500 | Gemini |
| 7 | PrEBM supplements | CC-4177 | Lonza |
| 8 | Charcoal stripped FBS | 12676029 | Gibco |
| 9 | T-PER buffer | 78510 | ThermoFisher |
| 10 | RIPA buffer | 89900 | ThermoFisher |
| 11 | pAKT-S473 antibody | 4060S | Cell Signaling |
| 12 | pAKT-T308 antibody | 2965S | Cell Signaling |
| 13 | Total AKT antibody | 4691S | Cell Signaling |
| 14 | Active β-catenin antibody | 8814S | Cell Signaling |
| 15 | H3K18lac antibody | PTM-1406RM | PTMbiolabs |
| 16 | GAPDH | 2118S | Cell Signaling |
| 17 | Liberase | 5401020001 | Sigma |
| 18 | CD45 antibody | 103138 | Biolegend |
| 19 | CD11b antibody | 101257 | Biolegend |
| 20 | CD11c antibody | 117339 | Biolegend |
| 21 | Ly6c antibody | 128006 | Biolegend |
| 22 | Ly6g antibody | 80-5931-U100 | Tonbo Biosciences |
| 23 | MHC-II antibody | 107631 | Biolegend |
| 24 | F4/80 antibody | 123147 | Biolegend |
| 25 | PD-1 antibody | 135228 | Biolegend |
| 26 | PD-L1 antibody | 25-5982-82 | Invitrogen |
| 27 | Ghost-viability dye | 13-0870-T100 | Tonbo Biosciences |
| 28 | CD3 antibody | 100216 | Biolegend |
| 29 | CD4 antibody | 100451 | Biolegend |
| 30 | CD8 antibody | 100748 | Biolegend |
| 31 | FoxP3 antibody | 126408 | Biolegend |
| 32 | Ki67 antibody | 652404 | Biolegend |
| 33 | CD206 antibody | 141708 | Biolegend |
| 34 | CD47 antibody | 127530 | Biolegend |
| 35 | Perm buffer | 421402 | Biolegend |
| 36 | Fix/perm buffer | 421401 | Biolegend |
| 37 | Annexin V-PI assay kit | 556547 | BD Biosciences |
| 38 | M-CSF | 315-02 | PeproTech |
| 39 | DMEM media | 17-205-CV | Corning |
| 40 | CTV dye | C34571 | ThermoFisher |
| 41 | Sodium-L-lactate | L7022 | Sigma |
| 42 | Lactate colorimetry kit | K627-100 | Biovision |
| 43 | Clodronate | SKU-CDL8909 | Encapsula Nano Sciences |
| 44 | Cytokine array | ARY028 | R&D systems |

## Notes

### Competing Interest Statement

The authors have declared no competing interest.

## References

1. H. Sung, J. Ferlay, R. L. Siegel, M. Laversanne, I. Soerjomataram, A. Jemal, F. Bray, Global Cancer Statistics 2020: GLOBOCAN Estimates of Incidence and Mortality Worldwide for 36 Cancers in 185 Countries. CA Cancer J Clin 71, 209–249 (2021).

2. T. S. Thomas, R. K. Pachynski, Treatment of Advanced Prostate Cancer. Mo Med 115, 156–161 (2018).

3. K. N. Chi, N. Agarwal, A. Bjartell, B. H. Chung, A. J. Pereira de Santana Gomes, R. Given, A. Juarez Soto, A. S. Merseburger, M. Ozguroglu, H. Uemura, D. Ye, K. Deprince, V. Naini, J. Li, S. Cheng, M. K. Yu, K. Zhang, J. S. Larsen, S. McCarthy, S. Chowdhury, T. Investigators, Apalutamide for Metastatic, Castration-Sensitive Prostate Cancer. N Engl J Med 381, 13–24 (2019).

4. M. R. Sydes, M. R. Spears, M. D. Mason, N. W. Clarke, D. P. Dearnaley, J. S. de Bono, G. Attard, S. Chowdhury, W. Cross, S. Gillessen, Z. I. Malik, R. Jones, C. C. Parker, A. W. S. Ritchie, J. M. Russell, R. Millman, D. Matheson, C. Amos, C. Gilson, A. Birtle, S. Brock, L. Capaldi, P. Chakraborti, A. Choudhury, L. Evans, D. Ford, J. Gale, S. Gibbs, D. C. Gilbert, R. Hughes, D. McLaren, J. F. Lester, A. Nikapota, J. O’Sullivan, O. Parikh, C. Peedell, A. Protheroe, S. M. Rudman, R. Shaffer, D. Sheehan, M. Simms, N. Srihari, R. Strebel, S. Sundar, S. Tolan, D. Tsang, M. Varughese, J. Wagstaff, M. K. B. Parmar, N. D. James, S. Investigators, Adding abiraterone or docetaxel to long-term hormone therapy for prostate cancer: directly randomised data from the STAMPEDE multi-arm, multi-stage platform protocol. Ann Oncol 29, 1235–1248 (2018).

5. C. J. Sweeney, Y. H. Chen, M. Carducci, G. Liu, D. F. Jarrard, M. Eisenberger, Y. N. Wong, N. Hahn, M. Kohli, M. M. Cooney, R. Dreicer, N. J. Vogelzang, J. Picus, D. Shevrin, M. Hussain, J. A. Garcia, R. S. DiPaola, Chemohormonal Therapy in Metastatic Hormone-Sensitive Prostate Cancer. N Engl J Med 373, 737–746 (2015).

6. I. D. Davis, A. J. Martin, M. R. Stockler, S. Begbie, K. N. Chi, S. Chowdhury, X. Coskinas, M. Frydenberg, W. E. Hague, L. G. Horvath, A. M. Joshua, N. J. Lawrence, G. Marx, J. McCaffrey, R. McDermott, M. McJannett, S. A. North, F. Parnis, W. Parulekar, D. W. Pook, M. N. Reaume, S. K. Sandhu, A. Tan, T. H. Tan, A. Thomson, E. Tu, F. Vera-Badillo, S. G. Williams, S. Yip, A. Y. Zhang, R. R. Zielinski, C. J. Sweeney, E. T. Investigators, A. the, U. New Zealand, G. Prostate Cancer Trials, Enzalutamide with Standard First-Line Therapy in Metastatic Prostate Cancer. N Engl J Med 381, 121–131 (2019).

7. M. R. Smith, M. Hussain, F. Saad, K. Fizazi, C. N. Sternberg, E. D. Crawford, E. Kopyltsov, C. H. Park, B. Alekseev, A. Montesa-Pino, D. Ye, F. Parnis, F. Cruz, T. L. J. Tammela, H. Suzuki, T. Utriainen, C. Fu, M. Uemura, M. J. Mendez-Vidal, B. L. Maughan, H. Joensuu, S. Thiele, R. Li, I. Kuss, B. Tombal, A. T. Investigators, Darolutamide and Survival in Metastatic, Hormone-Sensitive Prostate Cancer. N Engl J Med, (2022).

8. N. W. Clarke, A. Ali, F. C. Ingleby, A. Hoyle, C. L. Amos, G. Attard, C. D. Brawley, J. Calvert, S. Chowdhury, A. Cook, W. Cross, D. P. Dearnaley, H. Douis, D. Gilbert, S. Gillessen, R. J. Jones, R. E. Langley, A. MacNair, Z. Malik, M. D. Mason, D. Matheson, R. Millman, C. C. Parker, A. W. S. Ritchie, H. Rush, J. M. Russell, J. Brown, S. Beesley, A. Birtle, L. Capaldi, J. Gale, S. Gibbs, A. Lydon, A. Nikapota, A. Omlin, J. M. O’Sullivan, O. Parikh, A. Protheroe, S. Rudman, N. N. Srihari, M. Simms, J. S. Tanguay, S. Tolan, J. Wagstaff, J. Wallace, J. Wylie, A. Zarkar, M. R. Sydes, M. K. B. Parmar, N. D. James, Addition of docetaxel to hormonal therapy in low- and high-burden metastatic hormone sensitive prostate cancer: long-term survival results from the STAMPEDE trial. Ann Oncol 30, 1992–2003 (2019).

9. Y. Li, S. Malapati, Y. T. Lin, A. Patnaik, An integrative approach for sequencing therapies in metastatic prostate cancer. The American Journal of Hematology/Oncology 13, 26–31 (2017).

10. R. Montironi, A. Cimadamore, A. Lopez-Beltran, M. Scarpelli, G. Aurilio, M. Santoni, F. Massari, L. Cheng, Morphologic, Molecular and Clinical Features of Aggressive Variant Prostate Cancer. Cells 9, (2020).

11. V. Conteduca, S. Y. Ku, L. Fernandez, A. Dago-Rodriquez, J. Lee, A. Jendrisak, M. Slade, C. Gilbertson, J. Manohar, M. Sigouros, Y. Wang, R. Dittamore, R. Wenstrup, J. M. Mosquera, J. D. Schonhoft, H. Beltran, Circulating tumor cell heterogeneity in neuroendocrine prostate cancer by single cell copy number analysis. NPJ Precis Oncol 5, 76 (2021).

12. J. D. Wolchok, V. Chiarion-Sileni, R. Gonzalez, P. Rutkowski, J. J. Grob, C. L. Cowey, C. D. Lao, J. Wagstaff, D. Schadendorf, P. F. Ferrucci, M. Smylie, R. Dummer, A. Hill, D. Hogg, J. Haanen, M. S. Carlino, O. Bechter, M. Maio, I. Marquez-Rodas, M. Guidoboni, G. McArthur, C. Lebbe, P. A. Ascierto, G. V. Long, J. Cebon, J. Sosman, M. A. Postow, M. K. Callahan, D. Walker, L. Rollin, R. Bhore, F. S. Hodi, J. Larkin, Overall Survival with Combined Nivolumab and Ipilimumab in Advanced Melanoma. N Engl J Med 377, 1345–1356 (2017).

13. P. Sharma, J. P. Allison, Dissecting the mechanisms of immune checkpoint therapy. Nat Rev Immunol 20, 75–76 (2020).

14. M. Gamat, D. G. McNeel, Androgen deprivation and immunotherapy for the treatment of prostate cancer. Endocr Relat Cancer 24, T297–T310 (2017).

15. E. S. Antonarakis, J. M. Piulats, M. Gross-Goupil, J. Goh, K. Ojamaa, C. J. Hoimes, U. Vaishampayan, R. Berger, A. Sezer, T. Alanko, R. de Wit, C. Li, A. Omlin, G. Procopio, S. Fukasawa, K. I. Tabata, S. H. Park, S. Feyerabend, C. G. Drake, H. Wu, P. Qiu, J. Kim, C. Poehlein, J. S. de Bono, Pembrolizumab for Treatment-Refractory Metastatic Castration-Resistant Prostate Cancer: Multicohort, Open-Label Phase II KEYNOTE-199 Study. J Clin Oncol 38, 395–405 (2020).

16. P. Sharma, R. K. Pachynski, V. Narayan, A. Flechon, G. Gravis, M. D. Galsky, H. Mahammedi, A. Patnaik, S. Subudhi, M. Ciprotti, T. Duan, A. Saci, S. Hu, G. C. Han, K. Fizazi, Initial results from a phase II study of nivolumab (NIVO) plus ipilimumab (IPI) for the treatment of metastatic castration-resistant prostate cancer (mCRPC; CheckMate 650). Journal of Clinical Oncology 37, 142–142 (2019).

17. C. S. Jansen, N. Prokhnevska, H. T. Kissick, The requirement for immune infiltration and organization in the tumor microenvironment for successful immunotherapy in prostate cancer. Urol Oncol 37, 543–555 (2019).

18. D. Di Mitri, M. Mirenda, J. Vasilevska, A. Calcinotto, N. Delaleu, A. Revandkar, V. Gil, G. Boysen, M. Losa, S. Mosole, E. Pasquini, R. D’Antuono, M. Masetti, E. Zagato, G. Chiorino, P. Ostano, A. Rinaldi, L. Gnetti, M. Graupera, A. R. Martins Figueiredo Fonseca, R. Pereira Mestre, D. Waugh, S. Barry, J. De Bono, A. Alimonti, Re-education of Tumor-Associated Macrophages by CXCR2 Blockade Drives Senescence and Tumor Inhibition in Advanced Prostate Cancer. Cell Rep 28, 2156–2168 e2155 (2019).

19. Z. Lopez-Bujanda, C. G. Drake, Myeloid-derived cells in prostate cancer progression: phenotype and prospective therapies. J Leukoc Biol 102, 393–406 (2017).

20. N. Vitkin, S. Nersesian, D. R. Siemens, M. Koti, The Tumor Immune Contexture of Prostate Cancer. Front Immunol 10, 603 (2019).

21. A. J. Garcia, M. Ruscetti, T. L. Arenzana, L. M. Tran, D. Bianci-Frias, E. Sybert, S. J. Priceman, L. Wu, P. S. Nelson, S. T. Smale, H. Wu, Pten null prostate epithelium promotes localized myeloid-derived suppressor cell expansion and immune suppression during tumor initiation and progression. Mol Cell Biol 34, 2017–2028 (2014).

22. J. Singh, S. S. Sohal, A. Lim, H. Duncan, T. Thachil, P. De Ieso, Cytokines expression levels from tissue, plasma or serum as promising clinical biomarkers in adenocarcinoma of the prostate: a systematic review of recent findings. Ann Transl Med 7, 245 (2019).

23. J. D. Wu, L. M. Higgins, A. Steinle, D. Cosman, K. Haugk, S. R. Plymate, Prevalent expression of the immunostimulatory MHC class I chain-related molecule is counteracted by shedding in prostate cancer. J Clin Invest 114, 560–568 (2004).

24. T. Jamaspishvili, D. M. Berman, A. E. Ross, H. I. Scher, A. M. De Marzo, J. A. Squire, T. L. Lotan, Clinical implications of PTEN loss in prostate cancer. Nat Rev Urol 15, 222–234 (2018).

25. S. Wang, J. Gao, Q. Lei, N. Rozengurt, C. Pritchard, J. Jiao, G. V. Thomas, G. Li, P. Roy-Burman, P. S. Nelson, X. Liu, H. Wu, Prostate-specific deletion of the murine Pten tumor suppressor gene leads to metastatic prostate cancer. Cancer Cell 4, 209–221 (2003).

26. D. Robinson, E. M. Van Allen, Y. M. Wu, N. Schultz, R. J. Lonigro, J. M. Mosquera, B. Montgomery, M. E. Taplin, C. C. Pritchard, G. Attard, H. Beltran, W. Abida, R. K. Bradley, J. Vinson, X. Cao, P. Vats, L. P. Kunju, M. Hussain, F. Y. Feng, S. A. Tomlins, K. A. Cooney, D. C. Smith, C. Brennan, J. Siddiqui, R. Mehra, Y. Chen, D. E. Rathkopf, M. J. Morris, S. B. Solomon, J. C. Durack, V. E. Reuter, A. Gopalan, J. Gao, M. Loda, R. T. Lis, M. Bowden, S. P. Balk, G. Gaviola, C. Sougnez, M. Gupta, E. Y. Yu, E. A. Mostaghel, H. H. Cheng, H. Mulcahy, L. D. True, S. R. Plymate, H. Dvinge, R. Ferraldeschi, P. Flohr, S. Miranda, Z. Zafeiriou, N. Tunariu, J. Mateo, R. Perez-Lopez, F. Demichelis, B. D. Robinson, A. Sboner, M. Schiffman, D. M. Nanus, S. T. Tagawa, A. Sigaras, K. W. Eng, O. Elemento, A. Sboner, E. I. Heath, H. I. Scher, K. J. Pienta, P. Kantoff, J. S. de Bono, M. A. Rubin, P. S. Nelson, L. A. Garraway, C. L. Sawyers, A. M. Chinnaiyan, Integrative Clinical Genomics of Advanced Prostate Cancer. Cell 162, 454 (2015).

27. D. A. Fruman, H. Chiu, B. D. Hopkins, S. Bagrodia, L. C. Cantley, R. T. Abraham, The PI3K Pathway in Human Disease. Cell 170, 605–635 (2017).

28. V. B. Cetintas, N. N. Batada, Is there a causal link between PTEN deficient tumors and immunosuppressive tumor microenvironment? J Transl Med 18, 45 (2020).

29. W. Peng, J. Q. Chen, C. Liu, S. Malu, C. Creasy, M. T. Tetzlaff, C. Xu, J. A. McKenzie, C. Zhang, X. Liang, L. J. Williams, W. Deng, G. Chen, R. Mbofung, A. J. Lazar, C. A. Torres-Cabala, Z. A. Cooper, P. L. Chen, T. N. Tieu, S. Spranger, X. Yu, C. Bernatchez, M. A. Forget, C. Haymaker, R. Amaria, J. L. McQuade, I. C. Glitza, T. Cascone, H. S. Li, L. N. Kwong, T. P. Heffernan, J. Hu, R. L. Bassett, Jr., M. W. Bosenberg, S. E. Woodman, W. W. Overwijk, G. Lizee, J. Roszik, T. F. Gajewski, J. A. Wargo, J. E. Gershenwald, L. Radvanyi, M. A. Davies, P. Hwu, Loss of PTEN Promotes Resistance to T Cell-Mediated Immunotherapy. Cancer Discov 6, 202–216 (2016).

30. M. D. Wellenstein, K. E. de Visser, Cancer-Cell-Intrinsic Mechanisms Shaping the Tumor Immune Landscape. Immunity 48, 399–416 (2018).

31. A. Toso, A. Revandkar, D. Di Mitri, I. Guccini, M. Proietti, M. Sarti, S. Pinton, J. Zhang, M. Kalathur, G. Civenni, D. Jarrossay, E. Montani, C. Marini, R. Garcia-Escudero, E. Scanziani, F. Grassi, P. P. Pandolfi, C. V. Catapano, A. Alimonti, Enhancing chemotherapy efficacy in Pten-deficient prostate tumors by activating the senescence-associated antitumor immunity. Cell Rep 9, 75–89 (2014).

32. J. Blagih, F. Zani, P. Chakravarty, M. Hennequart, S. Pilley, S. Hobor, A. K. Hock, J. B. Walton, J. P. Morton, E. Gronroos, S. Mason, M. Yang, I. McNeish, C. Swanton, K. Blyth, K. H. Vousden, Cancer-Specific Loss of p53 Leads to a Modulation of Myeloid and T Cell Responses. Cell Rep 30, 481–496 e486 (2020).

33. C. Chen, Q. Cai, W. He, T. B. Lam, J. Lin, Y. Zhao, X. Chen, P. Gu, H. Huang, M. Xue, H. Liu, F. Su, J. Huang, J. Zheng, T. Lin, AP4 modulated by the PI3K/AKT pathway promotes prostate cancer proliferation and metastasis of prostate cancer via upregulating L-plastin. Cell Death Dis 8, e3060 (2017).

34. K. P. Hoeflich, M. Merchant, C. Orr, J. Chan, D. Den Otter, L. Berry, I. Kasman, H. Koeppen, K. Rice, N. Y. Yang, S. Engst, S. Johnston, L. S. Friedman, M. Belvin, Intermittent administration of MEK inhibitor GDC-0973 plus PI3K inhibitor GDC-0941 triggers robust apoptosis and tumor growth inhibition. Cancer Res 72, 210–219 (2012).

35. A. H. Awada, F. Morschhauser, J. P. Machiels, G. Salles, S. Rottey, S. Rule, D. Cunningham, F. Peyrade, C. Fruchart, H. T. Arkenau, I. Genvresse, K. Koechert, G. Cisternas, C. Granvil, C. Pena, L. Liu, PI3K inhibition and modulation of immune and tumor microenvironment markers by copanlisib in patients with non-Hodgkin’s lymphoma or advanced solid tumors. Annals of Oncology 29, viii20 (2018).

36. N. Liu, K. Haike, S. Glaeske, J. Paul, D. Mumberg, B. Kreft, K. Ziegelbauer, Copanlisib in combination with anti-PD-1 induces regression in animal tumor models insensitive or resistant to the monotherapies of PI3k and checkpoint inhibitors. Hematological Oncology 35, 257–258 (2017).

37. J. Yang, J. Nie, X. Ma, Y. Wei, Y. Peng, X. Wei, Targeting PI3K in cancer: mechanisms and advances in clinical trials. Mol Cancer 18, 26 (2019).

38. A. J. Armstrong, S. Halabi, P. Healy, J. J. Alumkal, C. Winters, J. Kephart, R. L. Bitting, C. Hobbs, C. F. Soleau, T. M. Beer, R. Slottke, K. Mundy, E. Y. Yu, D. J. George, Phase II trial of the PI3 kinase inhibitor buparlisib (BKM-120) with or without enzalutamide in men with metastatic castration resistant prostate cancer. Eur J Cancer 81, 228–236 (2017).

39. C. Sweeney, S. Bracarda, C. N. Sternberg, K. N. Chi, D. Olmos, S. Sandhu, C. Massard, N. Matsubara, B. Alekseev, F. Parnis, V. Atduev, G. L. Buchschacher, Jr., R. Gafanov, L. Corrales, M. Borre, D. Stroyakovskiy, G. V. Alves, E. Bournakis, J. Puente, M. L. Harle-Yge, J. Gallo, G. Chen, J. Hanover, M. J. Wongchenko, J. Garcia, J. S. de Bono, Ipatasertib plus abiraterone and prednisolone in metastatic castration-resistant prostate cancer (IPATential150): a multicentre, randomised, double-blind, phase 3 trial. Lancet 398, 131–142 (2021).

40. C. Yong, D. L. Moose, N. Bannick, W. R. Gutierrez, M. Vanneste, R. Svensson, P. Breheny, J. A. Brown, R. D. Dodd, M. B. Cohen, M. D. Henry, Locally invasive, castrate-resistant prostate cancer in a Pten/Trp53 double knockout mouse model of prostate cancer monitored with non-invasive bioluminescent imaging. PLoS One 15, e0232807 (2020).

41. Y. C. Shen, A. Ghasemzadeh, C. M. Kochel, T. R. Nirschl, B. J. Francica, Z. A. Lopez-Bujanda, M. A. Carrera Haro, A. Tam, R. A. Anders, M. J. Selby, A. J. Korman, C. G. Drake, Combining intratumoral Treg depletion with androgen deprivation therapy (ADT): preclinical activity in the Myc-CaP model. Prostate Cancer Prostatic Dis 21, 113–125 (2018).

42. J. S. de Bono, U. De Giorgi, D. N. Rodrigues, C. Massard, S. Bracarda, A. Font, J. A. Arranz Arija, K. C. Shih, G. D. Radavoi, N. Xu, W. Y. Chan, H. Ma, S. Gendreau, R. Riisnaes, P. H. Patel, D. J. Maslyar, V. Jinga, Randomized Phase II Study Evaluating Akt Blockade with Ipatasertib, in Combination with Abiraterone, in Patients with Metastatic Prostate Cancer with and without PTEN Loss. Clin Cancer Res 25, 928–936 (2019).

43. P. Martin, Y. N. Liu, R. Pierce, W. Abou-Kheir, O. Casey, V. Seng, D. Camacho, R. M. Simpson, K. Kelly, Prostate epithelial Pten/TP53 loss leads to transformation of multipotential progenitors and epithelial to mesenchymal transition. Am J Pathol 179, 422–435 (2011).

44. M. Feng, W. Jiang, B. Y. S. Kim, C. C. Zhang, Y.-X. Fu, I. L. Weissman, Phagocytosis checkpoints as new targets for cancer immunotherapy. Nature Reviews Cancer 19, 568–586 (2019).

45. R. L. Elstrom, D. E. Bauer, M. Buzzai, R. Karnauskas, M. H. Harris, D. R. Plas, H. Zhuang, R. M. Cinalli, A. Alavi, C. M. Rudin, C. B. Thompson, Akt stimulates aerobic glycolysis in cancer cells. Cancer Res 64, 3892–3899 (2004).

46. D. Zhang, Z. Tang, H. Huang, G. Zhou, C. Cui, Y. Weng, W. Liu, S. Kim, S. Lee, M. Perez-Neut, J. Ding, D. Czyz, R. Hu, Z. Ye, M. He, Y. G. Zheng, H. A. Shuman, L. Dai, B. Ren, R. G. Roeder, L. Becker, Y. Zhao, Metabolic regulation of gene expression by histone lactylation. Nature 574, 575–580 (2019).

47. T. Vidotto, C. M. Melo, E. Castelli, M. Koti, R. B. Dos Reis, J. A. Squire, Emerging role of PTEN loss in evasion of the immune response to tumours. Br J Cancer 122, 1732–1743 (2020).

48. N. Liu, J. Luo, D. Kuang, S. Xu, Y. Duan, Y. Xia, Z. Wei, X. Xie, B. Yin, F. Chen, S. Luo, H. Liu, J. Wang, K. Jiang, F. Gong, Z. H. Tang, X. Cheng, H. Li, Z. Li, A. Laurence, G. Wang, X. P. Yang, Lactate inhibits ATP6V0d2 expression in tumor-associated macrophages to promote HIF-2alpha-mediated tumor progression. J Clin Invest 129, 631–646 (2019).

49. A. Viola, F. Munari, R. Sanchez-Rodriguez, T. Scolaro, A. Castegna, The Metabolic Signature of Macrophage Responses. Front Immunol 10, 1462 (2019).

50. J. Chen, X. Cao, B. Li, Z. Zhao, S. Chen, S. W. T. Lai, S. A. Muend, G. K. Nossa, L. Wang, W. Guo, J. Ye, P. P. Lee, M. Feng, Warburg Effect Is a Cancer Immune Evasion Mechanism Against Macrophage Immunosurveillance. Front Immunol 11, 621757 (2020).

51. X. Geeraerts, J. Fernandez-Garcia, F. J. Hartmann, K. E. de Goede, L. Martens, Y. Elkrim, A. Debraekeleer, B. Stijlemans, A. Vandekeere, G. Rinaldi, R. De Rycke, M. Planque, D. Broekaert, E. Meinster, E. Clappaert, P. Bardet, A. Murgaski, C. Gysemans, F. A. Nana, Y. Saeys, S. C. Bendall, D. Laoui, J. Van den Bossche, S. M. Fendt, J. A. Van Ginderachter, Macrophages are metabolically heterogeneous within the tumor microenvironment. Cell Rep 37, 110171 (2021).

52. D. Kim, J. Wang, S. B. Willingham, R. Martin, G. Wernig, I. L. Weissman, Anti-CD47 antibodies promote phagocytosis and inhibit the growth of human myeloma cells. Leukemia 26, 2538–2545 (2012).

53. A. A. Barkal, K. Weiskopf, K. S. Kao, S. R. Gordon, B. Rosental, Y. Y. Yiu, B. M. George, M. Markovic, N. G. Ring, J. M. Tsai, K. M. McKenna, P. Y. Ho, R. Z. Cheng, J. Y. Chen, L. J. Barkal, A. M. Ring, I. L. Weissman, R. L. Maute, Engagement of MHC class I by the inhibitory receptor LILRB1 suppresses macrophages and is a target of cancer immunotherapy. Nat Immunol 19, 76–84 (2018).

54. A. A. Barkal, R. E. Brewer, M. Markovic, M. Kowarsky, S. A. Barkal, B. W. Zaro, V. Krishnan, J. Hatakeyama, O. Dorigo, L. J. Barkal, I. L. Weissman, CD24 signalling through macrophage Siglec-10 is a target for cancer immunotherapy. Nature 572, 392–396 (2019).

55. Y. P. Chen, H. J. Kim, H. Wu, T. Price-Troska, J. C. Villasboas, S. Jalali, A. L. Feldman, A. J. Novak, Z. Z. Yang, S. M. Ansell, SIRPalpha expression delineates subsets of intratumoral monocyte/macrophages with different functional and prognostic impact in follicular lymphoma. Blood Cancer J 9, 84 (2019).

56. B. Wang, T. Tian, K. H. Kalland, X. Ke, Y. Qu, Targeting Wnt/beta-Catenin Signaling for Cancer Immunotherapy. Trends Pharmacol Sci 39, 648–658 (2018).

57. S. A. Haney, High-Content Screening Approaches That Minimize Confounding Factors in RNAi, CRISPR, and Small Molecule Screening. Methods Mol Biol 1683, 113–130 (2018).

58. C. Massard, K. N. Chi, D. Castellano, J. de Bono, G. Gravis, L. Dirix, J. P. Machiels, A. Mita, B. Mellado, S. Turri, J. Maier, D. Csonka, A. Chakravartty, K. Fizazi, Phase Ib dose-finding study of abiraterone acetate plus buparlisib (BKM120) or dactolisib (BEZ235) in patients with castration-resistant prostate cancer. Eur J Cancer 76, 36–44 (2017).

59. B. S. Carver, C. Chapinski, J. Wongvipat, H. Hieronymus, Y. Chen, S. Chandarlapaty, V. K. Arora, C. Le, J. Koutcher, H. Scher, P. T. Scardino, N. Rosen, C. L. Sawyers, Reciprocal feedback regulation of PI3K and androgen receptor signaling in PTEN-deficient prostate cancer. Cancer Cell 19, 575–586 (2011).

60. P. H. Cha, J. H. Hwang, D. K. Kwak, E. Koh, K. S. Kim, K. Y. Choi, APC loss induces Warburg effect via increased PKM2 transcription in colorectal cancer. Br J Cancer 124, 634–644 (2021).

61. Y. L. Park, H. P. Kim, Y. W. Cho, D. W. Min, S. K. Cheon, Y. J. Lim, S. H. Song, S. J. Kim, S. W. Han, K. J. Park, T. Y. Kim, Activation of WNT/beta-catenin signaling results in resistance to a dual PI3K/mTOR inhibitor in colorectal cancer cells harboring PIK3CA mutations. Int J Cancer 144, 389–401 (2019).

62. Y. N. Liu, W. Abou-Kheir, J. J. Yin, L. Fang, P. Hynes, O. Casey, D. Hu, Y. Wan, V. Seng, H. Sheppard-Tillman, P. Martin, K. Kelly, Critical and reciprocal regulation of KLF4 and SLUG in transforming growth factor beta-initiated prostate cancer epithelial-mesenchymal transition. Mol Cell Biol 32, 941–953 (2012).

63. X. Y. Zhao, P. J. Malloy, A. V. Krishnan, S. Swami, N. M. Navone, D. M. Peehl, D. Feldman, Glucocorticoids can promote androgen-independent growth of prostate cancer cells through a mutated androgen receptor. Nat Med 6, 703–706 (2000).

64. Y. Kfoury, N. Baryawno, N. Severe, S. Mei, K. Gustafsson, T. Hirz, T. Brouse, E. W. Scadden, A. A. Igolkina, K. Kokkaliaris, B. D. Choi, N. Barkas, M. A. Randolph, J. H. Shin, P. J. Saylor, D. T. Scadden, D. B. Sykes, P. V. Kharchenko, C. as part of the Boston Bone Metastases, Human prostate cancer bone metastases have an actionable immunosuppressive microenvironment. Cancer Cell 39, 1464–1478 e1468 (2021).

65. R. Zappasodi, I. Serganova, I. J. Cohen, M. Maeda, M. Shindo, Y. Senbabaoglu, M. J. Watson, A. Leftin, R. Maniyar, S. Verma, M. Lubin, M. Ko, M. M. Mane, H. Zhong, C. Liu, A. Ghosh, M. Abu-Akeel, E. Ackerstaff, J. A. Koutcher, P. C. Ho, G. M. Delgoffe, R. Blasberg, J. D. Wolchok, T. Merghoub, CTLA-4 blockade drives loss of Treg stability in glycolysis-low tumours. Nature 591, 652–658 (2021).

66. S. Butenko, S. K. Satyanarayanan, S. Assi, S. Schif-Zuck, N. Sher, A. Ariel, Transcriptomic Analysis of Monocyte-Derived Non-Phagocytic Macrophages Favors a Role in Limiting Tissue Repair and Fibrosis. Front Immunol 11, 405 (2020).

67. F. O. Martinez, S. Gordon, M. Locati, A. Mantovani, Transcriptional profiling of the human monocyte-to-macrophage differentiation and polarization: new molecules and patterns of gene expression. J Immunol 177, 7303–7311 (2006).

68. M. Martin, Cutadapt removes adapter sequences from high-throughput sequencing reads. 2011 17, 3 (2011).

69. A. Dobin, C. A. Davis, F. Schlesinger, J. Drenkow, C. Zaleski, S. Jha, P. Batut, M. Chaisson, T. R. Gingeras, STAR: ultrafast universal RNA-seq aligner. Bioinformatics 29, 15–21 (2013).

70. M. I. Love, W. Huber, S. Anders, Moderated estimation of fold change and dispersion for RNA-seq data with DESeq2. Genome Biol 15, 550 (2014).

71. D. Croft, G. O’Kelly, G. Wu, R. Haw, M. Gillespie, L. Matthews, M. Caudy, P. Garapati, G. Gopinath, B. Jassal, S. Jupe, I. Kalatskaya, S. Mahajan, B. May, N. Ndegwa, E. Schmidt, V. Shamovsky, C. Yung, E. Birney, H. Hermjakob, P. D’Eustachio, L. Stein, Reactome: a database of reactions, pathways and biological processes. Nucleic Acids Res 39, D691–697 (2011).

